# A multi-tiered map of EMT defines major transition points and identifies vulnerabilities

**DOI:** 10.1101/2021.06.01.446492

**Authors:** Indranil Paul, Dante Bolzan, Ahmed Youssef, Keith A. Gagnon, Heather Hook, Gopal Karemore, Michael UJ Oliphant, Weiwei Lin, Qian Liu, Sadhna Phanse, Carl White, Dzmitry Padhorny, Sergei Kotelnikov, Guillaume P. Andrieu, Christopher S. Chen, Pingzhao Hu, Gerald V. Denis, Dima Kozakov, Brian Raught, Trevor Siggers, Stefan Wuchty, Senthil K. Muthuswamy, Andrew Emili

## Abstract

Epithelial to mesenchymal transition (EMT) is a complex cellular program proceeding through a hybrid E/M state linked to cancer-associated stemness, migration and chemoresistance. Deeper molecular understanding of this dynamic physiological landscape is needed to define events which regulate the transition and entry into and exit from the E/M state. Here, we quantified >60,000 molecules across ten time points and twelve omic layers in human mammary epithelial cells undergoing TGFβ-induced EMT. Deep proteomic profiles of whole cells, nuclei, extracellular vesicles, secretome, membrane and phosphoproteome defined state-specific signatures and major transition points. Parallel metabolomics showed metabolic reprogramming preceded changes in other layers, while single-cell RNA sequencing identified transcription factors controlling entry into E/M. Covariance analysis exposed unexpected discordance between the molecular layers. Integrative causal modeling revealed co-dependencies governing entry into E/M that were verified experimentally using combinatorial inhibition. Overall, this dataset provides an unprecedented resource on TGFβ signaling, EMT and cancer.

## Introduction

Epithelial to mesenchymal transition (EMT) regulates cell plasticity during embryonic development, wound healing, fibrosis and cancer, where polarized epithelial (E) cells dedifferentiate, transition through intermediate hybrid states (E/M) and acquire mesenchymal (M) properties (Nieto et al., 2016). In cancer, cells in E/M state possess several clinically important attributes of circulating tumor cells (CTCs) and are responsible for EMT-associated stemness, chemoresistance, immune evasion and metastasis (Dongre and Weinberg, 2019). Complete molecular characterization of E/M states, and the mechanisms driving plasticity between E®E/M®M transitions will enable development of refined mechanistic models and discovery of new therapeutic strategies.

Approximately 150 genes are currently described as hallmarks of EMT (MSigDB database) that were identified from studies measuring the expression of ‘endpoint’ markers (e.g. CDH1, MUC1, VIM, FN1) to track the process (Sha et al., 2019). Although EMT is frequently studied as a transcriptionally-driven program (Yang et al., 2020), the poor correlation between genomic alterations or mRNA levels and proteins in tumors (Liu et al., 2016) highlights the need for multi-level analysis. Furthermore, EMT is likely an *emergent phenomenon*, where the shifts in cell physiology and phenotype are orchestrated by intra- and extra-cellular signaling (Sigston and Williams, 2017), extensive receptor-ligand crosstalk, protein relocalization (Hung and Link, 2011) and metabolic adaptations (Thomson et al., 2019) that requires an integrated multi-omic approach. However, most attempts made to model EMT primarily invoked gene-regulatory networks (GRNs) controlled by key transcription factors (TFs) and miRNAs. These models are usually based on a restricted number of factors and thus do not capture the multi-layered architecture of signaling during EMT (Hong et al., 2015; Zhang et al., 2014). To date, no EMT-focused studies have simultaneously measured metabolite and gene expression changes at different functional levels (e.g., mRNA, total protein, nucleus, secretome, etc.).

Consequentially, several aspects of EMT remain unclear. This includes dependencies between molecular layers, secreted molecules, and specific signatures at various stages of EMT, kinetics and scope of metabolic reprogramming, dynamics of subcellular protein localizations and ligand-receptor mediated intercellular crosstalk. To bridge these gaps, we employed multiple high-throughput platforms including microarray, scRNAseq and precision mass spectrometry (MS) to quantify molecules spanning 12 distinct layers of biological information. Our ability to integrate this information allowed us to define signatures of E/M states and identify molecular regulators of key transition points.

## RESULTS

### A comprehensive resource on TGFβ-induced EMT

The human mammary epithelial cell line MCF10A is widely used to study TGFβ (transforming growth factor β) induced EMT. To generate temporal expression maps of evolving EMT landscape, cells were treated with TGFβ (TGF-β1; 10 ng/mL) over 12 days (0, 4 hrs, 1–6, 8 & 12 days), spanning multiple ‘omics’ layers and employing complementary technologies (**Fig. 1A-B, Fig. S1A-C**). After stringent quality control (see **STAR Methods**), we report nanoLC-MS/MS-based quantifications (**Fig. 1C, Table S1**) of 6,540 whole cell (*WC*), 4,198 nuclear (*Nuc*), 2,223 plasma membrane (*Mem*), 1,209 extracellular vesicle (*EV*) and 1,133 secreted (*Sec*) proteins. Using a serial enrichment workflow (see **STAR Methods**) we also quantified the total phosphoproteome (*Phos*; 8,741 high-confidence sites on 2,254 proteins; including 6,975 Ser, 962 Thr and 140 Tyr residues), N-glycoproteome (*Glyco*; 549 proteins), acetylome (*Acet*; 349 sites on 165 proteins) and peptidome (*Pep*; 547 peptides from 202 proteins). For proteomics, samples for ten time point were multiplexed using isobaric tandem mass tags (TMT-10), enabling higher throughput and robust comparisons. In addition to proteins, we tracked cellular metabolism in the same samples by nanoLC-MS/MS-based untargeted metabolomics, quantifying 4,259 HMDB-indexed endogenous small molecules (*Metabol*). Furthermore, we measured 23,787 gene transcripts (mRNA) and 2,578 microRNAs (miRNA) using microarrays. To assess cellular heterogeneity, we employed scRNAseq to quantify transcriptomes of 1,913 individual cells (>200 cells per time point) undergoing EMT. In total, this study provides temporal quantifications of >60,000 proteins, phosphosites, mRNAs, miRNAs, and metabolites combined, in addition to 9,785 mRNAs in scRNAseq dataset (**Fig. 1C**).

**Figure 1.**
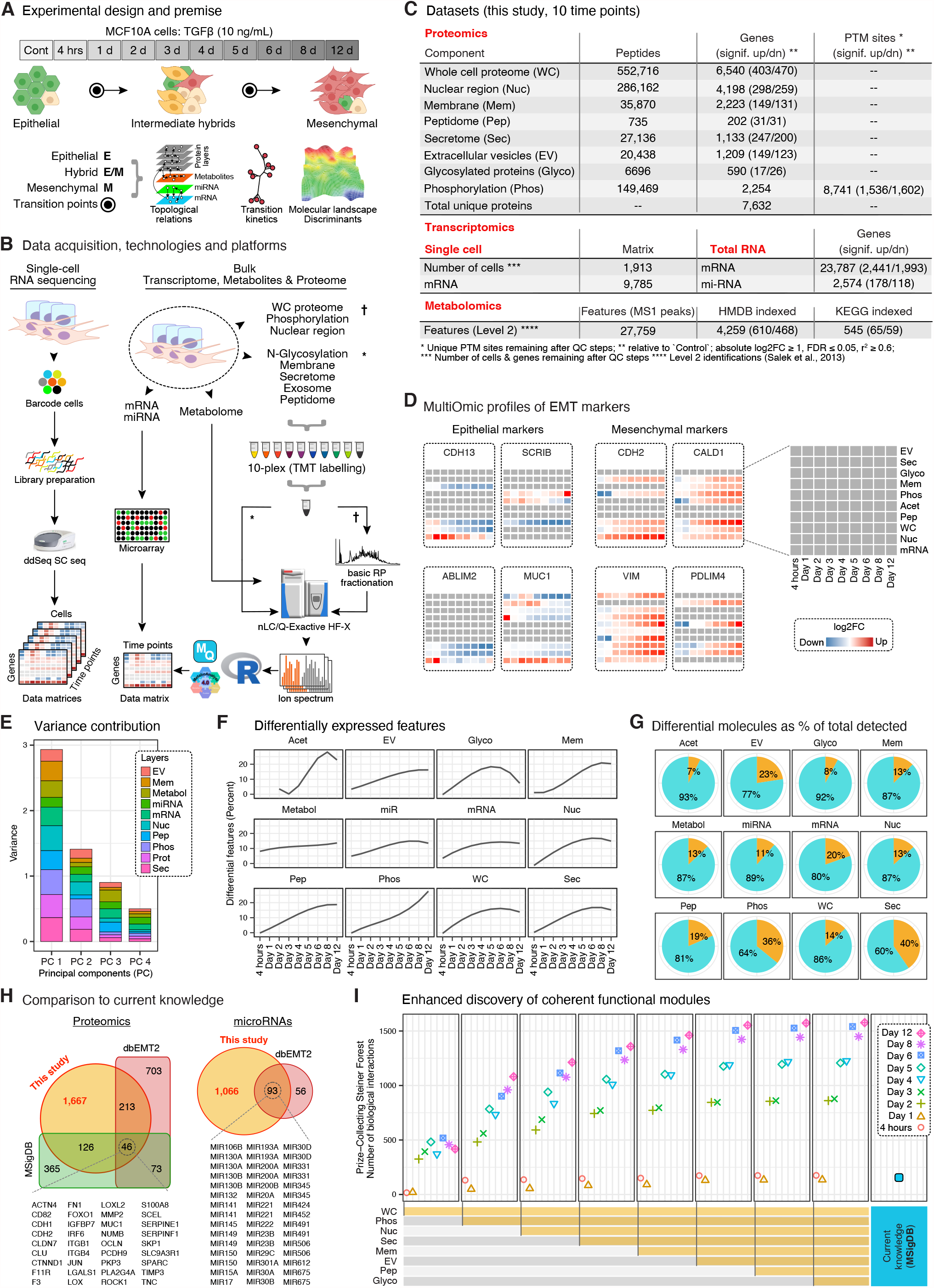
A multi-dimensional resource on TGFβ-induced EMT. (A) MCF10A cells exposed to TGFβ for indicated time points were used to study the molecular landscapes during EMT. (B) Samples from 3 biological replicates were aliquoted and multiple technologies were employed to quantify various molecular layers. (C) An overview of numbers of molecules quantified in various layers. (D) Expression snapshots of some well-known EMT markers. Heatmaps show log2FC values (relative to Control, adj. *p-value* < 0.05). (E) Variance explained by each layer over 4 principal components. (F) Ridge plot showing differential molecules, as % of total quantified, for each layer. (G) Pie chart showing the overall fraction (in %) of differential molecules (yellow portion) relative to all molecules quantified in each layer. (H) Overlap between established EMT databases (MSigDB, www.gsea-msigdb.org; dbEMT2.0, http://dbemt.bioinfo-minzhao.org) and differential proteins and miRNAs (adj. *p-value* < 0.05; r^2^ ≥ 0.6 & |log2FC| ≥ 1) from this study. (I) Differential molecules (proteins, miRNAs, metabolites) were used to assess the number of coherent functional modules, i.e., known interactions between molecules, by employing the Prize-Collecting Steiner Forest algorithm on a network compiled from PathwayCommons, miRTarBase and STITCH.

Subcellular enrichments were performed using previously established MS-compatible protocols (see **STAR Methods**), yielding high purity as determined through keyword matching against a cellular compartment annotation database (**Fig. S1D**). Quantitative reproducibility across the 3 biological replicates was excellent (**Fig. S1E**). The expression profiles in **Fig. 1D** provide a snapshot of concurrent changes of a given gene over various layers during EMT. We reproduced expected expression behavior for many established markers of EMT, including an increase of M markers VIM, CDH2 and concomitant decrease of E markers SCRIB, MUC1.

Using strict criteria for differential expression (Benjamini-Hochberg adj. *p-value* < 0.05; r^2^ ≥ 0.6 & |log2FC| ≥ 1) we identified >10,000 significantly regulated molecules (**Table S2**). We found that molecular abundances were highly variable across layers and time points (**Fig. S1F**) and all layers contributed significantly to the overall variation in the system (**Fig. 1E**). While each layer showed substantive alterations, the magnitudes (**Fig. S1G**), response profiles (**Fig. 1F**) and fraction of differentially expressed molecules (**Fig. 1G**) of each omic data set during the time course varied significantly. These findings provide direct evidence for global reorganization of cell physiology during EMT. For example, *Phos* and *Sec* showed the highest fractional change (36% & 40%, respectively) (**Fig. 1G**), suggesting extensive intra- and extra-cellular signaling during EMT (Scheel et al., 2011). As such, this study is the largest experimental description of EMT till date and adds new depth to our current understanding of TGFβ signaling and EMT (**Fig. 1H, I**).

### Integrative multi-tiered topology of EMT reveals key transition points and identifies molecular drivers

Although E®M progression is considered a continuum, the existence of E, hybrid E/M, and M states have been reported for MCF10A cells (Zhang et al., 2014), and cancer tissues (Liu et al., 2019; Pastushenko et al., 2018). Using a phylogenetic clustering approach (Hughes and Friedman, 2009) we estimated ‘distances’ between time steps to understand the transition kinetics (**Fig. 2A**). We observed that up to 24 hours cells maintained their parental E type, while day 2 marked a swift exit from E, and entry into E/M which continued through Day 5, after which cells gradually entered the M state. Cells in day 2 to 5 can be further sub-divided into E/M–1 (Day 2/3, late E) and E/M–2 (Day 4/5, early M) states. These cellular reconfigurations agreed with principal component analysis of individual layers, such as *Mem, Phos* and *WC* (**Fig. S2A**).

**Figure 2.**
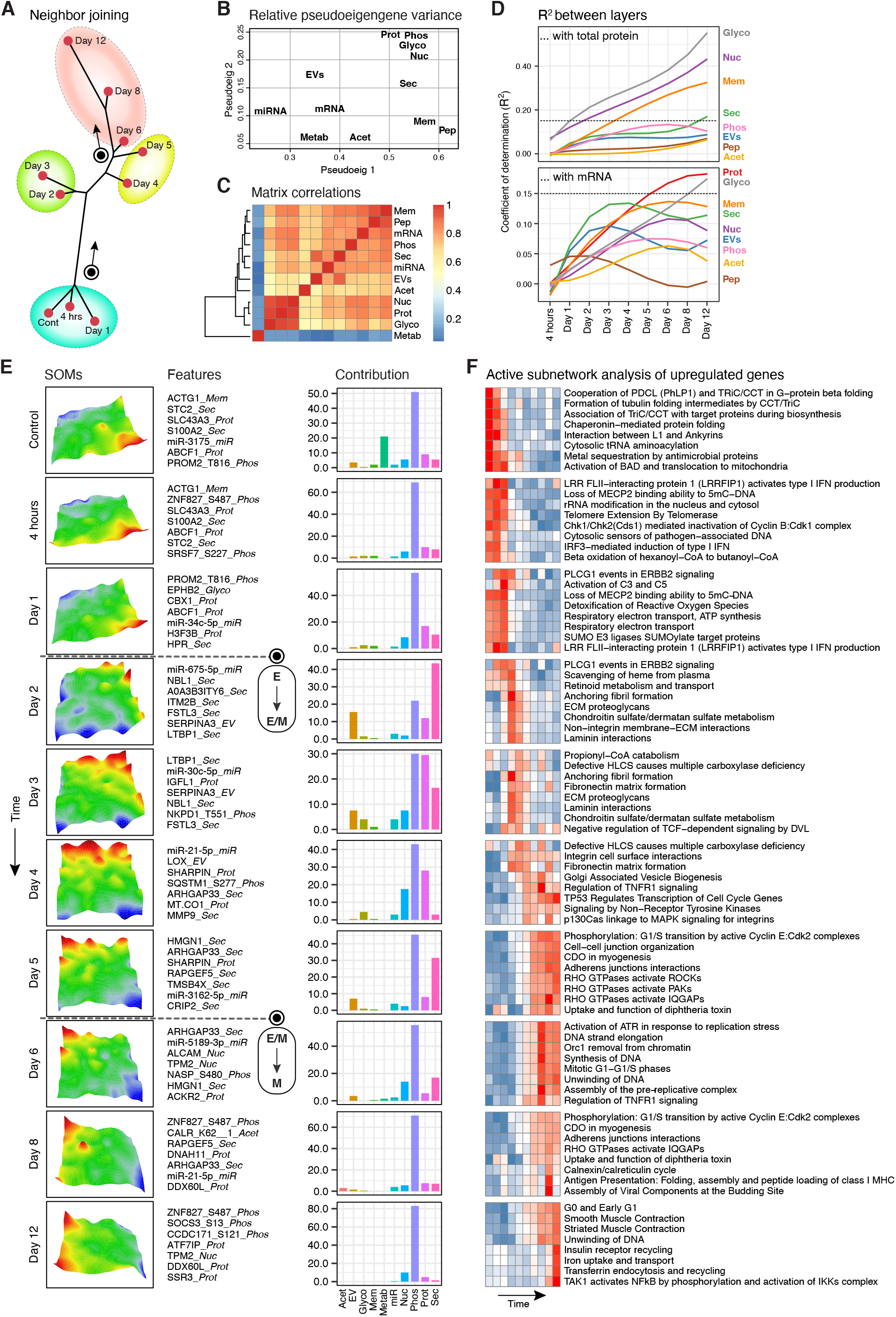
The topological architecture of EMT. (A) Phylogenetic neighbor-joining tree reveals similarities (=distances) between time points. (B) Combined pseudo-eigenvalues space of all datasets, indicating the contribution of each dataset to the eigenvalue (variance). (C) Matrix correlations between each pair of datasets. (D) Line plots show the distribution of adjusted coefficient of determination (*R*^*2*^) values between layers as a function of time points. (E) *Left panel*. SOM portraits. Color gradient refers to over-or under-expression of metagenes in each time-point compared to the mean expression level of the metagene in the pool of all time points: red = high, yellow/green = intermediate levels and blue = low (see **STAR Methods** for details). *Middle panel*. Representative examples that appeared among the highest-ranking features (top 1%) in each time-point. *Right panel*. Barplot shows the number of features that were contributed by each layer among the highest-ranking features (top 1%) in time-point. (F) Heatmap depicts sample-wise pathway scores, which were derived from enrichment analysis of active subnetworks using the highest-ranking molecules (top 1%) for each SOM.

Our observations suggest a distinct and complementary role for each molecular layer in shaping the transition (**Fig 2B, Fig 1E-G, Fig S1F, G**). Although, the topology was generally correlated (**Fig. 2C**), pairwise coefficients of determination (adj. *R*^*2*^) revealed an unexpectedly low concordance between total proteins and other proteomic layers (mean *R*^*2*^ ranging from 0.015 for *Acet* to 0.299 for *Glyco*) (**Fig. 2D, Upper panel**). The discordance was even stronger between mRNA and various proteomic layers (mean *R*^*2*^ ranging from 0.011 for *Pep* to just 0.109 for *WC*) (**Fig. 2D, Lower panel**), which increased further with EMT progression. Collectively, these observations illustrate that at a systems level mRNA quantity is a poor proxy of protein abundance. Even total protein quantity is an unreliable predictor of post-translational modifications or subcellular trafficking, which ultimately determines signaling output.

Next, we modeled the regulatory patterns of molecules using a 3-step computational workflow. First, we used a regression strategy for time-course measurements (Conesa et al., 2006) to remove residual noise (time dimension) and non-reliable (replicates) components. Second, an unsupervised machine learning approach (self-organizing maps, SOMs) (Wirth et al., 2012) was applied on the retained molecules to generate time-point specific SOM portraits (**Fig. 2E, Fig. S2B-D**, see **STAR Methods** for details). SOMs are a powerful integration tool for diverse global datasets to extract underlying patterns of co-regulation (Tamayo et al., 1999). Third, we performed pathway enrichments of these SOMs to elucidate the overarching functional themes at each time-step of EMT (**Fig. 2F**).

SOMs traced the temporal unfolding of the transition and provided molecular fingerprints of each time-step. Molecules ranking high (top 1% i.e., rank ≤ 250 & log2FC ≥ 1) with SOMs for control to day 1 (430 unique molecules) primarily included proteins characteristic of an E state (**Fig. 2E, Table S3**), such as, ACTG1 (*Mem*; establishes cell junctions and shape) and EPHB2 (*Glyco*; regulates angiogenesis, contact dependent adhesion and migration; tumor suppressor). SOMs for days 6–12 (252 unique molecules) contained known M markers, e.g., CALD1, CDH2, FLNA, FN1, FSTL1, LGALS1, NT5E, TAGLN, TPM1, TPM2, TPM4 and VIM. This group also included molecules with established roles in cancer and EMT, including miR-5189 (*miR*; targets ARF6 which internalizes CDH1), CALR (*Acet*; Lys 62; calcium homeostasis, promotes metastasis, imparts resistance to anoikis), ALCAM (*Nuc*; prognostic marker in multiple cancers linked to nuclear translocation of β-catenin and stemness), ARHGAP33 (*Sec*; regulates intracellular trafficking), RAPGEF5 (*Sec*; promotes nuclear translocation of β-catenin) and SOCS3 (*Phos*; Thr 3/13; E3 ligase, E3 ligase inhibiting TGFβ signaling). As suggested in **Fig 2A**, the first major shift, E→E/M, occurred at day 2 and was likely driven by molecules peaking at its corresponding SOM (95 molecules) such as miR-675-5p/H19 (*miR*; induces HIF1α, SNAIL activity), LTBP1 (*Sec*; master regulator of integrin-dependent TGFβ activation) and CD44 (*Sec*; signal transduction). Notably, many of the molecules identified in the SOM analysis, to our knowledge, have not been previously linked to TGFβ signaling or EMT, presumably because they do not show clear changes in *mRNA* or *WC* but in ‘other’ molecular layers, such as *EV* or *Sec* (**Fig. S2E**). This contrasts with most MSigDB hallmarks of EMT for which we captured clear transcriptional profiles (**Fig. S2F**).

Gene-set enrichment analysis using active subnetworks which yields more robust inferences than traditional approaches (Ulgen et al., 2019), identified 237 significant pathways (**Fig. 2F, Table S3**), discretized across sequential steps of EMT. For example, ‘Beta oxidation of hexanoyl-CoA to butanoyl-CoA’ declined as cells leave E and enter E/M, indicating reprogramming of mitochondrial fatty acid β-oxidation, consistent with a metastatic phenotype (Ma et al., 2018). Conversely, ‘RHO GTPases mediated activation of ROCKs/PAKs/IQGAPs’ increased as cells leave E/M and enter M, suggestive of their key role at this stage of EMT (Ungefroren et al., 2018). The E/M specifically were associated with migration-associated pathways such as ‘anchoring fibril formation’, ‘ECM proteoglycans’ and ‘laminin interactions’, consistent with their shared property with CTCs.

Overall, we catalogued complex kinetics of thousands of molecules spanning multiple molecular layers during EMT. Importantly, we identify critical transition points during EMT and predict signatures specific to each stage, e.g., E/M, with potential clinical value.

### Metabolomics reveals kinetics and predicts novel enzyme-metabolite associations

TGFβ regulates Warburg effect in cancer cells, but may also regulate other metabolic pathways with implications for cancer management (Hua et al., 2019). Active subnetwork analysis of *WC* and *Phos* datasets identified 13 enriched KEGG metabolic pathways during TGFβ-induced EMT (**Fig. S3A**). Notably, significant enrichment (*p–value* ≤ 0.05) for any pathway was only observed after day 2 and included processes such as steroid hormone biosynthesis (SHB), sphingolipid metabolism and (SLM) glycosaminoglycan degradation (GMGD). To evaluate these gene-centric inferences of metabolic phenotypes, we directly profiled intra-cellular small molecules by applying an optimized untargeted metabolomics workflow to the same set of samples. Using stringent criteria (see **STAR Methods**), we quantified >4,000 putative HMDB-compounds covering a wide range of chemical classes (**Fig. 3A**). Using phylogenetic clustering (**Fig. 3B**) and SOM analysis (**Fig. 3C**) driven solely by the *Metabol* dataset, we observed the E→E/M transition as early as 4 hours, followed by another transition after day 1. This indicates rapid modulation of cellular metabolism by TGFβ, preceding changes in most other layers. The E/M→M transition occurred around day 5, in line with the integrative analysis.

**Figure 3.**
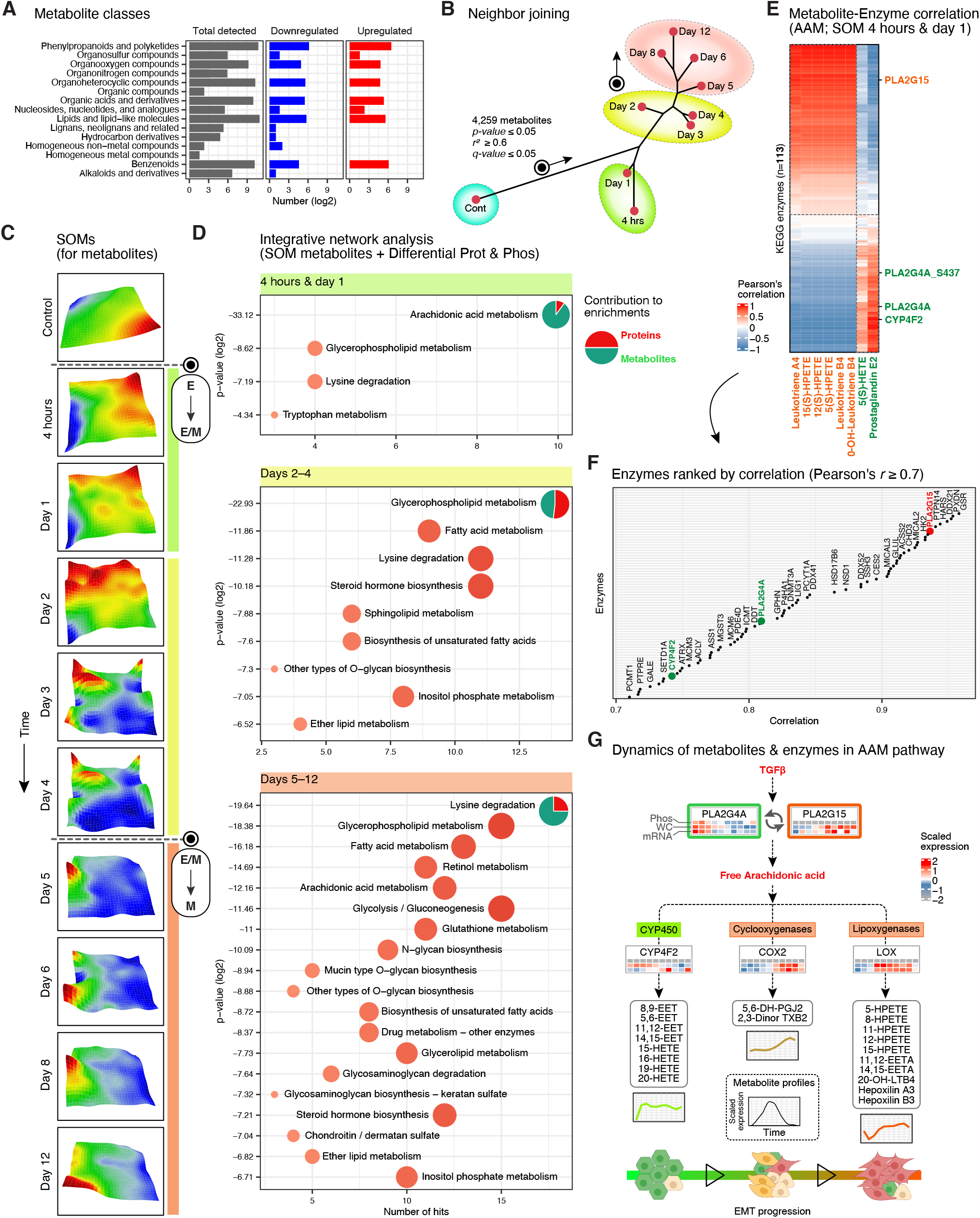
Dynamics of TGFβ-induced metabolic adaptations. (A) Barplot showing the ‘class’ distribution of quantified metabolite features in HMBD. Also shown are the relative proportions of differentially expressed features for each class (*p-value* ≤ 0.01, absolute log2FC ≥ 1). (B) Neighbor-joining tree for the *Metabol* dataset. (C) SOM analysis of the *Metabol* dataset. (D) Top 10% of metabolite features of each SOM were grouped based on clusters in ‘B’ and used for “Network analysis” using MetaboAnalyst (https://www.metaboanalyst.ca). (E) Identified metabolites of AAM pathway in SOM for 4 hours to day 1 were taken and Pearson’s correlation computed with known metabolic enzymes (KEGG) quantified in the *Phos & WC* datasets. (F) The plot shows enzymes ranked according to their Pearson’s correlation with metabolites of AAM pathway, as detected in this study. (G) Schematics of information flow from TGFβ signaling to AAM pathway, mediated by known enzymes. Heatmaps and line plots display ‘standardized’ expression values.

To glean further insights, we performed integrative network analysis of metabolite SOMs with differential molecules in *WC* and *Phos* using MetaboAnalyst (Chong et al., 2019) (**Fig. 3D**). This analysis reiterated several enriched pathways predicted with protein expression alone, e.g., SHB, SLM, GMGD, glycerophospholipid metabolism (GPLM) and lysine degradation (LD) (**Fig. S3A**). However, integration of metabolite and protein data within the framework of metabolite SOMs revealed pathway activities representative of key transition steps of EMT driven primarily by corresponding metabolite signatures (**Fig. S3B-D**). Indeed, we observed that arachidonic acid metabolism (AAM), GPLM and LD pathways were activated within 4 hours of TGFβ stimulation, which was not captured by gene-set analysis. Consistent with observations in **Fig 2F**, processes such as fatty acid metabolism, SHB and SLM appear after day 2, as cells prepare for a metastatic phenotype (Koundouros and Poulogiannis, 2020).

Pairwise-correlation based integration of metabolite profiles with proteomics measurements from the same samples could enable mechanistic predictions and aid discovery of novel players. To explore this, we chose AAM as an example. Arachidonic acid is an omega-6 fatty acid stored as membrane phosphoglycerolipid. Its cytosolic release enables stoichiometric chain reactions and results in >100 functionally diverse compounds (Hanna and Hafez, 2018) impacting processes such as redox state, proliferation, apoptosis and chemotaxis (Tallima and El Ridi, 2018). We computed Pearson’s correlations between abundance of KEGG-annotated enzymes, in *WC* and *Phos*, and metabolites of AAM pathway, in *Metabol*, quantified at SOM for 4 hours to day 1 (**Fig. 3E, F**). Overall, we found that metabolites mapping to AAM either rapidly increased and then stabilized (Cytochrome P450, CYP450, branch) or showed a delayed but consistent increase over the time course (Cyclooxygenase, COX, and Lipoxygenase, LOX, branches) (**Fig. 3G**), suggesting fine-tuned regulation of the different branches. Encouragingly, our enzyme-metabolite association map identified PLA2G4A, the rate-limiting phospholipase of AAM pathway (Hanna and Hafez, 2018), among the top candidates (**Fig. 3F**). Interestingly, correlation and expression profiles of PLA2G15, another phospholipase, (**Fig. 3E-G**) indicated a potential enzyme-mediated switch from CYP450 to COX/LOX branches during E→E/M.

Overall, these observations reveal the kinetics of metabolic reprogramming during TGFβ induced EMT. We found metabolites and protein signatures coordinating processes such as AAM, GPLM and LD during key stages of the transition. We also demonstrate how our enzyme-metabolite correlation map could be used as a resource to predict novel enzymes for observed metabolic changes.

### scRNAseq analysis reveals heterogeneous responses to TGFβ and novel transcriptional regulators of EMT

scRNAseq studies in murine epithelial Py2T cells treated with TGFβ (Krishnaswamy et al., 2018), MCF10A cells undergoing confluence-dependent EMT (McFaline-Figueroa et al., 2019) and LPS-induced EMT in alveolar epithelial cells (Riemondy et al., 2019), provided valuable insights into EMT. However, a temporal analysis of TGFβ-induced EMT to understand the transition states has not been reported.

After quality control, we retained 1,913 single cells with a combined depth of 9,785 genes (**Fig. S4A, Table S1**). As anticipated, many of the top expressing genes (TGFB1, TPT1, KRT6A, TMSB10, MT2A) are key players in EMT (**Fig. S4B**). Interestingly, similar to many primary human tumors (Puram et al., 2018), we did not observe an explicit loss of several classical E markers at the transcript level (**Fig. 4A, Fig. S4C**), suggesting post-transcriptional regulation, a strategy which could be energetically economical for tumor cells (Lambert et al., 2017).

**Figure 4.**
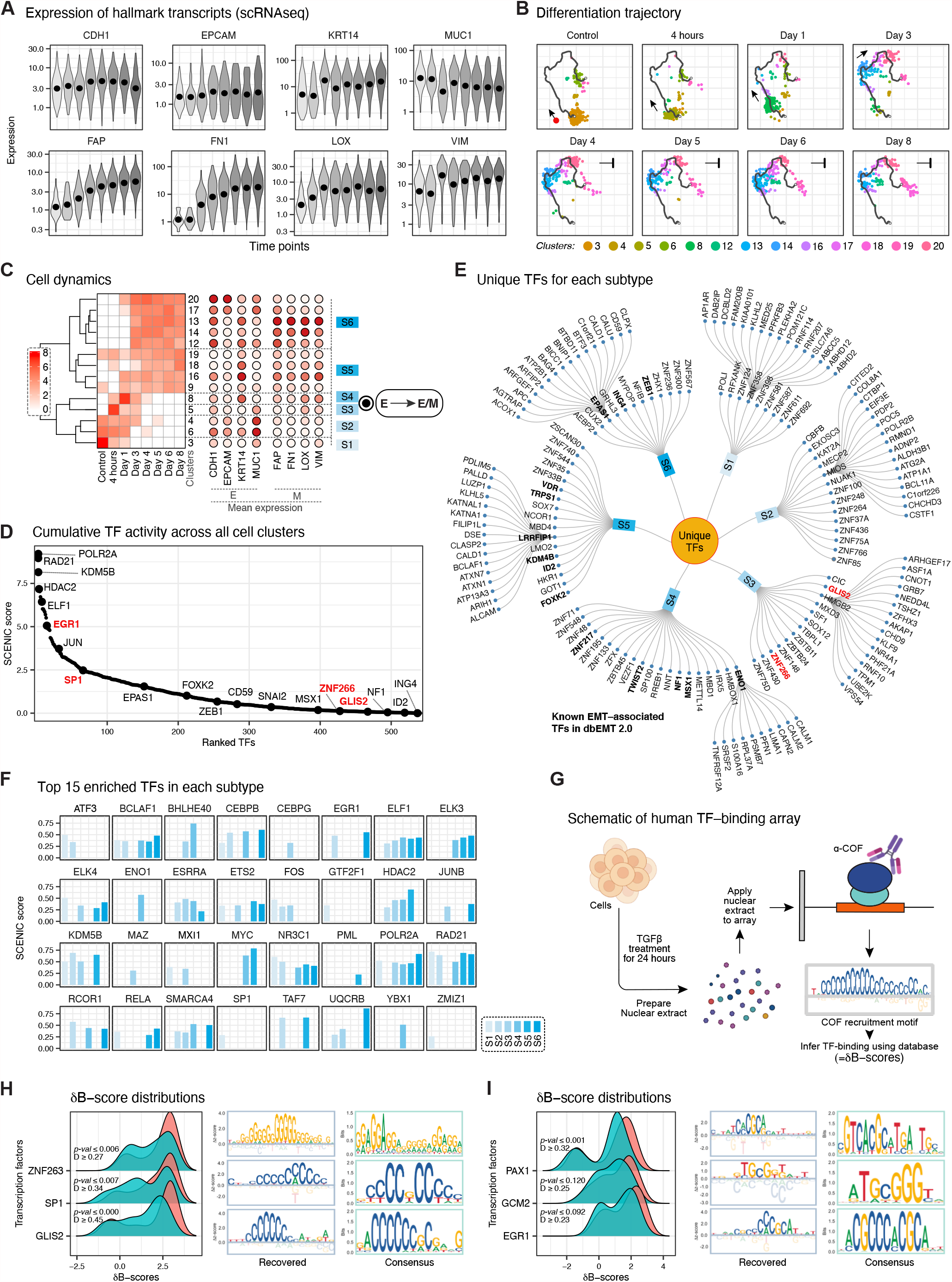
scRNAseq analysis reveals cellular dynamics and novel TFs for EMT. (A) Violin plots showing expression of well-known EMT hallmarks in each time point. (B) UMAP of scRNAseq by Monocle3. Dots represent single cells and are colored by inferred clusters, while trajectories depict cells during EMT. (C) Heatmap showing the number of cells (in log2 scale) in each cluster of partition P2. Mean expression of some well-known E and M markers in each cluster are also shown. (D) The plot shows all TFs ranked according to their SCENIC score. TF names are shown for the 5 TFs with the highest scores and some well-known EMT associated TFs. (E) Tree displays unique TFs identified by SCENIC for each subtype. TFs highlighted in ‘bold’ are known players in EMT. The genes in the outer circle are representative examples used by SCENIC to infer TF activity. (F) Barplot shows the inferred activities of top 15 TFs based on SCENIC scores in each subtype. (G) Schematics of the human TF-binding array workflow (see **STAR Methods** for details). (H) & (I) Density plots of ∂B-scores of indicated TFs. Two-sided Kolmogorov-Smirnov test was performed to evaluate the significance of distribution differences between Control and TGFβ treated conditions.

To understand the stages of cell differentiation, we took advantage of the multiple time points in our dataset (as opposed to a ‘pseudo-time’), and identified 20 cell clusters in 3 disjoint partitions, using Monocle3 (**Fig. 4B, Fig. S4D**). Partition P2 (12 clusters) represents the primary EMT axis while P1/P3 predominantly expressed genes related to cell cycle (**Fig. S4E**) and were ignored for further analysis. We observed that C3 responded strongly to TGFβ (**Fig. 4C**), C4/6 resisted EMT, C5/C8 are the ‘transition’ states and C13/14 represented terminal M cells (in terms of hallmark M markers; **Fig. 4C, right panel**). Examining clusters C9/18/19 which were composed of cells from nearly all time points, suggests presence of stable M-type cells in MCF10A populations and appears to be at transcriptional impasse for TGFβ signaling. Notably, using scRNAseq, we could observe an E→E/M transition, but the E/M→M transition, as revealed by integrative analysis, was not clear.

To explore the underlying gene expression program, we used hierarchical clustering to group the individual clusters into 6 subtypes (**Fig. 4C**) and employed SCENIC (Aibar et al., 2017) to infer TFs and GRNs underlying these subtypes (**Fig. D-F; Table S5**). For each subtype, we identified several unique (**Fig. 4E**) or highly active TFs (**Fig. 4F**), including both established and novel players. Several TFs implicated in EMT (TWIST2, FOXK2, ZEB1, ID2, MSX1, ING4) were over-represented in S4-S6, which corresponds to later stages of EMT. In contrast, direct evidence of mechanistic links of TFs enriched in early stages of EMT (i.e., S3, S4) are lacking, indicating gaps in current EMT models. Using human TF–binding arrays (see **STAR Methods, Fig. 4G**) we confirmed elevated activity of three S3 TFs, GLIS2, SP1 and ZNF266 upon TGFβ induction (**Fig. 4H**). In addition to providing experimental evidence to our predictions, the TF-binding array also revealed several other TFs potentially playing important roles at the early stages of EMT (**Fig. 4I**).

Overall, we provide a high-resolution temporal map of gene expression programs of individual cells as they respond to TGFβ signaling and undergo EMT. More generally, our data suggests that most transcriptional changes occur at early time points (also **Fig. 2D, Lower panel**), followed by further adaptations driven predominantly by post-transcriptional mechanisms.

### Spatial regulation of proteins and inter-cellular communication during EMT

Regulation of protein distribution is a crucial signaling mechanism (Ferrell, 1998), but remains insufficiently understood in EMT. The average Pearson’s correlation between proteins quantified in multiple cellular compartments (CCs) ranged between *r* = 0.12 to 0.58 indicating fine-tuned regulation of protein distributions (**Fig. 5A**). We found 3,965 proteins localizing to ≥2 CCs (**Fig. S5A**), which we categorized into 2 classes as follows: Class I proteins (1,424) displayed a correlated trend (*r* ≥ 0.4) consistently across all CC pairs suggesting regulation primarily at or before the translation step (**Fig. 5B, C; Table S5**). Class II proteins (1,205) displayed anti-correlated trend (*r* ≤ –0.4) between any two CCs, implying active post-translational control of their asymmetric distributions (**Fig. 5D, E; Table S5**).

**Figure 5.**
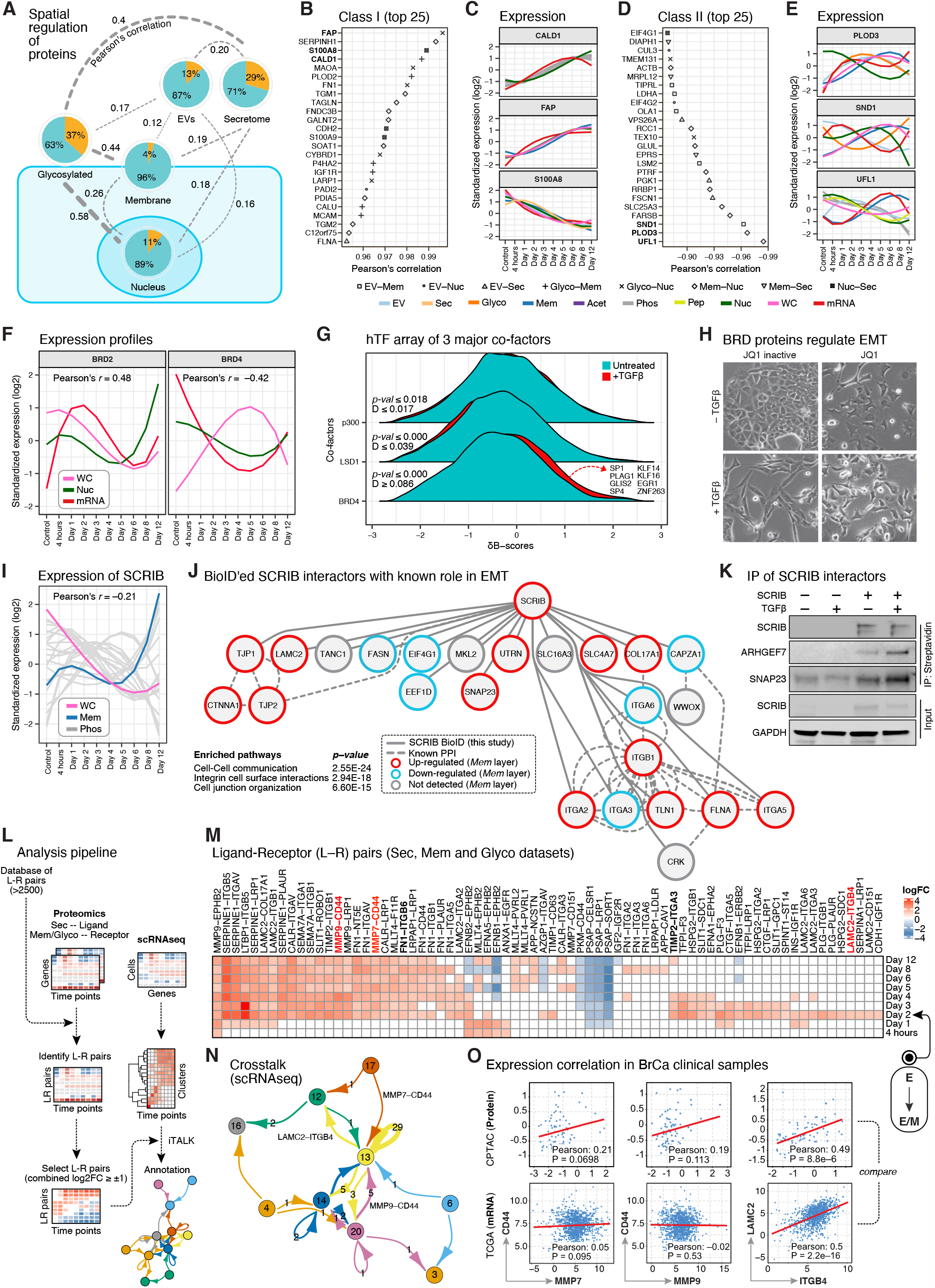
Spatial regulation of proteins and intercellular communication. (A) The schematic summarizes Pearson’s coefficients between overlapping proteins of the indicated layers. Each pie chart depicts the fraction of differential proteins (orange slice) with respect to all proteins quantified in the layer. (B) The plot displays top 25 Class I proteins. Each pair is represented by a different shape. (C) Expression profiles of top 3 Class I proteins. Each colored line represents a molecular layer. (D) The plot displays top 25 Class II proteins. Each pair is represented by a different shape. (E) Expression profiles of top 3 Class II proteins. Each colored line represents a molecular layer. (F) Expression profiles of BRD2 and BRD4 in various layers. Legend as in E. (G) Density plots of ∂B-scores of indicated co-factors. Two-sided Kolmogorov-Smirnov test was performed to evaluate the significance of difference in distributions. (H) Phase-contrast images after 6 days of TGFβ treatment in presence or absence of active JQ-1 (100 nM) or its inactive analogue. (I) Expression profile of SCRIB in various layers. Legend as in E. (J) PPI network of SCRIB interactors identified using BioID. (K) Immunoblots showing interactions of a few SCRIB partners identified using BioID. (L) Schematics of analysis pipeline for discovering active L-R pairs (see **STAR Methods**). (M) Heatmap showing combined log2FCs of L-R pairs in the *Sec* and *Mem/Glyco* datasets. (N) Network plot showing L-R interactions detected between different P2 cell clusters. (O) Scatter plot of correlations between indicated gene-pairs in Breast invasive carcinoma samples. Regression line is shown in red.

The Bromodomain and ExtraTerminal (BET) cofactors, BRD2 and BRD4, were identified as Class II proteins (**Fig. 5F**). Using TF-binding arrays, we verified enhanced global recruitment of BRD4, but not two other cofactors, p300 and LSD1, (**Fig. 5G**) indicating its specific role during EMT. Indeed, treatment with a selective BET inhibitor JQ-1 suppressed EMT (**Fig. 5H**). Another notable Class II protein was SCRIB (**Fig. 5I**), which regulates apical-basal polarity and directional migration by acting as a molecular scaffold through protein-protein interactions (PPIs) (Bonello and Peifer, 2019). Using an *in* vivo proximity ligation (BioID) screen of SCRIB ((**Fig. S5B**, see **STAR Methods**), we identified multiple novel interactors (**Table S6**) of which many have known roles in EMT (**Fig. 5J**). Using immunoprecipitation, we verified interactions of SCRIB with SNAP23 and ARHGEF7 (**Fig. 5K**).

Subtypes within a cell population can differ in their capacity to send and receive signals, with implications for metastasis and drug resistance (Kim et al., 2018; Tabassum and Polyak, 2015). To map the inter-cellular communication between subgroups of cells during TGFβ-induced EMT, we integrated proteomics and scRNAseq data to perform a systems-wide survey of ligand-receptor (L-R) pair mediated crosstalk (**Fig. 5L**). First, using a database of >2,500 curated binary L-R interactions (Ramilowski et al., 2015), we searched for pairs of L and R in our *Sec* and *Mem/Glyco* datasets, respectively, assuming that co-directional expression changes in L and/or R of a pair (FDR adj. *p-value* < 0.05 and combined L-R |log2FC| ≥1) can indicate biological role. Currently, at least two L-R pairs are implicated in TGFβ-induced EMT (Heldin et al., 2012). Our analysis detected 67 upregulated and 12 downregulated L-R pairs at any given time point following TGFβ treatment (**Fig. 5M, Table S7**). Notably, none of these pairs have been directly implicated in TGFβ signaling or EMT, although individually many of the identified L or R occur frequently in the context of EMT and/or cancer. For instance, LAMC2 with its 7 receptors (CD151, COL17A1, ITGA2, ITGA3, ITGA6, ITGB1, ITGB4) exhibit significant alteration during EMT. LAMC2 is overexpressed in cancers (Garg et al., 2014) and its silencing can reverse EMT (Pei et al., 2019). The cognate receptors, CD151, ITGA3, ITGA6 and ITGB1 synergize with TGFβ signaling to promote metastatic behavior (Pellinen et al., 2018; Sadej et al., 2010; Shirakihara et al., 2013; Zhang et al., 2017).

Next, by systematically comparing the expression patterns of L & R (identified above) among the 16 clusters (identified in our scRNAseq dataset), we obtained cell-cell communication networks (sender → receiver) (**Fig. 5N, Fig. S5C, Table S7**). For example, C13 cells, which appeared at day 3 and showed highest M genes expression (**Fig. 4C**), produced the receptor CD44 to its cognate ligand MMP7 expressed by C17 (**Fig. S5D**). Together, this suggests that communication between cell subgroups (C17:MMP7 → C13:CD44) may exist during EMT which might have potential ramifications for tumor growth. Interestingly, our analysis suggests a global switch in cell surface proximal signaling cascades at day 2, corresponding to E→E/M transition, and likely modulating processes characteristic of E/M cells, e.g., migration (e.g., FN1–ITGB6) and stemness (e.g., TIMP2–ITGA3).

As independent corroboration, several identified L-R pairs showed strong correlation (**Fig. 5O**) in human breast invasive carcinoma samples (Cancer Genome Atlas Network, 2012; Cerami et al., 2012). Notably, a slightly stronger correlation between the L–R pairs was observed with CPTAC proteomics data than with TCGA mRNA datasets.

Our study provides a comprehensive analysis of TGFβ-triggered subcellular trafficking as cells undergo EMT. Such translocations, potentially driven by differential PPI, could mediate the tight coordination between functional modules (e.g., SCRIB complex) and EMT phenotypes such as cytoskeletal rearrangement. We also uncovered novel cell-cell communication pathways via L-R interactions in driving EMT, representing an untapped clinical opportunity.

### Modeling phosphoproteome dynamics during EMT reveals kinase susceptibilities

Phosphoregulatory mechanisms are a key aspect of TGFβ signaling and EMT. We confidently quantified 8,741 phosphosites (p-sites; 6,975 Ser, 962 Thr and 140 Tyr residues) (**Fig. S6A-C**) over a dynamic range of 10^6^ orders of magnitude (**Fig. S6D**), and phospho–STY frequencies (**Fig. S6E**) in line with previous reports (D’Souza et al., 2014) mapping to 2,254 proteins (**Fig. S6F**). Of all p-sites, 3,138 (35.8%) were differentially regulated in at least one time point (**Fig. S6G**). At protein level, different patterns of regulation were noted; some proteins, such as DEK, VIM and MISP, were regulated at ∼90% of detected sites, some proteins, such as CAV1, CAMK2 and GOLGA1, showed a ∼50% mixture of regulated and unregulated sites, while others such as AHNAK, PML and BCLAF1 showed ∼2% differential sites (**Fig. 6A**). Interestingly, the fraction of regulated p-sites in several proteins, e.g., VIM, increased with EMT progression (**Fig. 6B**).

**Figure 6.**
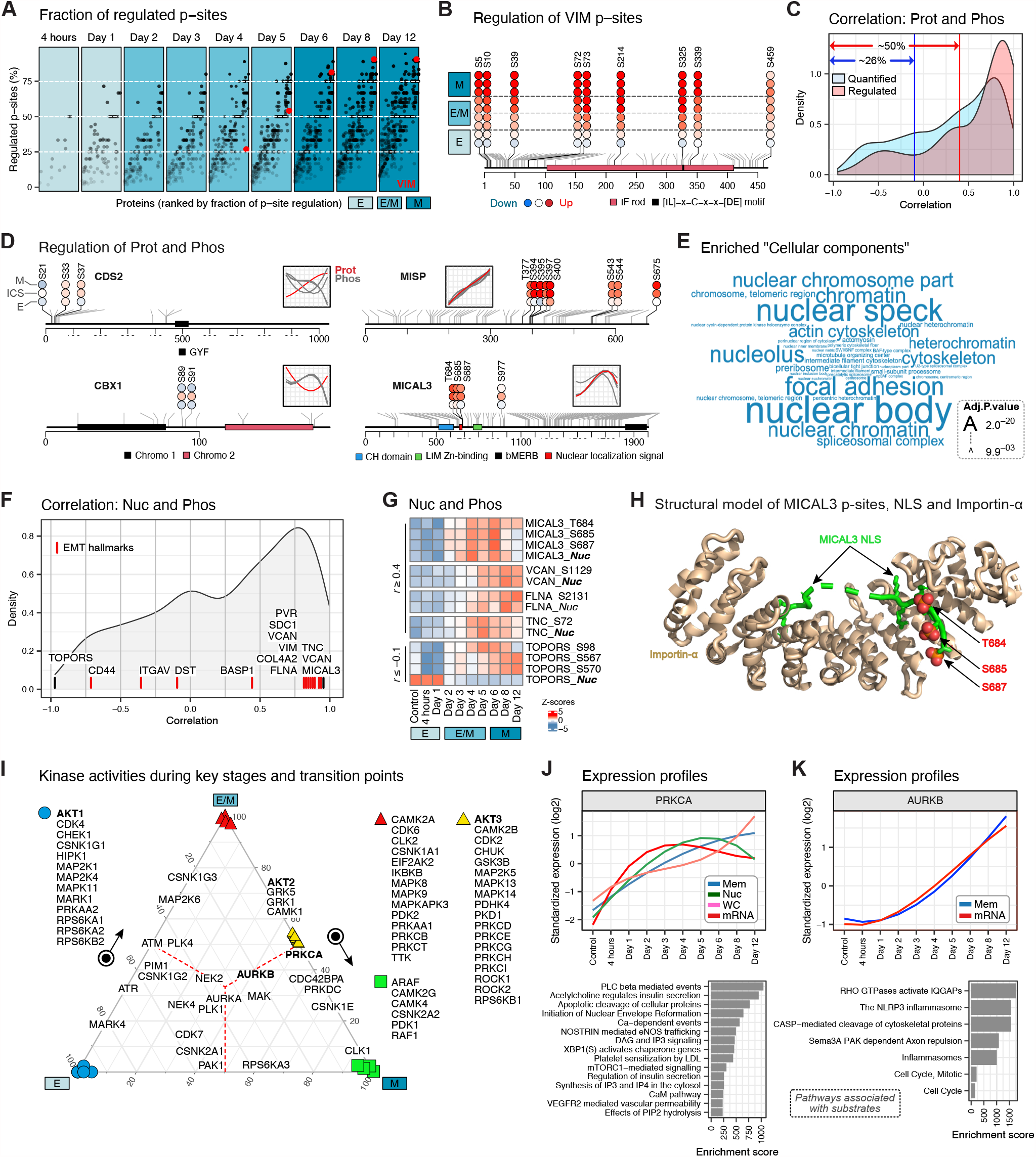
Phosphoproteome dynamics during EMT. (A) Fraction of detected p-sites on a protein that are regulated during EMT. (B) Detected p-sites on VIM and their expression during EMT. The gray lines indicate all p-sites that are catalogued in PhosphoSitePlus database. (C) Distribution of Pearson’s correlation between expression of proteins and p-sites detected on them. (D) Schematic showing two examples each where expression of proteins and p-sites showed either low (≤–0.1; CDS2, CBX1) or high (≥0.4; MISP, MICAL3) correlation. (E) Gene ontology enrichment of genes with at least a single regulated p-site at any time point. (F) Distribution of Pearson’s correlation between expression of proteins detected in *Nuc* layer and the p-sites detected on them in the *Phos* layer. A few EMT hallmarks are highlighted. (G) Heatmaps of expression profiles of indicated molecules in *Nuc* and *Phos* layers. (H) Structural model of MICAL3 p-sites, NLS and Importin-α. (I) Ternary plot of kinase activity scores binned into 3 broad stages of EMT, i.e., E, ICS, and M. (J) and (K) *Top*. Expression of PRKCA or AURKB as detected in various layers. *Bottom*. Pathway enrichment of all differential substrates of PRKCA or AURKB detected in our dataset.

We observed that in ∼50%, p-sites dynamics were not explained (*r* ≤ 0.4) by a corresponding change at the protein level (**Fig. 6C, D**). Interestingly, for ∼26%, a directionally opposite change between phosphorylation and the corresponding protein abundance was noted (*r* ≤ –0.1), suggesting effects on protein stability. Phospho-regulated proteins were enriched for ‘nucleus’, ‘cytoskeleton’ and ‘focal adhesion’ annotations (**Fig. 6E**) reflecting the importance of CC remodeling during EMT.

We also computed correlations between nuclear localization profiles (*Nuc*) and phosphorylation kinetics (*Phos*) of individual proteins (**Fig. 6F, G**). For instance, phosphorylation of MICAL3 at T684 and S685 regulates CSCs by promoting symmetric division (Tominaga et al., 2019). The pattern of phosphorylation of MICAL3 at residues T684, S685 and S687 (**Fig. 6D**, MICAL3) which are located at the consensus NLS motif suggests a role in regulating nuclear translocation of MICAL3 at E→E/M transition. Indeed, the bipartite NLS motif of MICAL3 interacts with Importin-α and the p-sites T684, S685, S687 are directly adjacent to the binding interface (**Fig. 6H**).

To analyze stage-specific kinase activities we generated a ternary model which distinguishes active kinases into 3 broad stages (i.e., E, E/M, and M) of EMT (**Fig. 6I**). An example that illustrates the utility of this model are AKT isoforms that have distinct and opposing roles during cancer development (Hinz and Jücker, 2019). We predict AKT1 is strongly associated with the E-stage, which is consistent with its role in maintaining the E phenotype (Li et al., 2016). In fact, depletion of AKT1 in MCF10A cells promoted TGFβ-induced E→E/M transition (Iliopoulos et al., 2009). Our model further predicts key roles for AKT2 and AKT3 at E/M→M transition. Indeed, AKT2 and AKT3 were associated with tumor invasiveness, stemness and sensitivity to drug treatment (Chin et al., 2014), key characteristics of the E/M populations (Dongre and Weinberg, 2019). Among several other kinases (**Fig. 6J, Fig. S6H**), our ternary model predicted key roles for PRKCA and AURKB at the junction of E→E/M and E/M→M transitions. PRKCA is reportedly a hub and therapeutic target for EMT-induced breast CSCs (Tam et al., 2013). Similarly, inhibition of AURKB was found to reverse EMT and reduce breast cancer metastasis *in vivo* (Zhang et al., 2020).

Overall, we reveal the rich intricacies of the phosphoregulome during EMT, identified functional p-sites, predict novel kinase susceptibilities, and provide a mechanistic framework to enhance understanding of the signaling mechanisms during EMT.

### Integrative systems causal model of EMT identifies mechanistic vulnerabilities

Systems biology approaches that combine multiple molecular types (proteins, mRNAs, miRNAs, metabolites) into a framework of established knowledge allow for a rich assessment of a biological context (Hawe et al., 2019). Using experimentally validated functional priors (compiled from ENCODE, PhosphoSitePlus, SignaLink 2.0, SIGNOR 2.0, HINT, miRTarBase and MetaBridge), we combined causal inference and PCSF (Prize Collecting Steiner Forest) (Akhmedov et al., 2017) to construct hierarchical mechanistic models of the EMT program (**Fig. 7A**, see **STAR Methods**). The final ‘EMT network’ comprised of 3,255 edges connecting 2,217 molecules, including 723 kinase/phosphatase–substrate, 1,407 TF–target, 746 miRNA–target and 31 metabolite–gene interactions.

**Figure 7.**
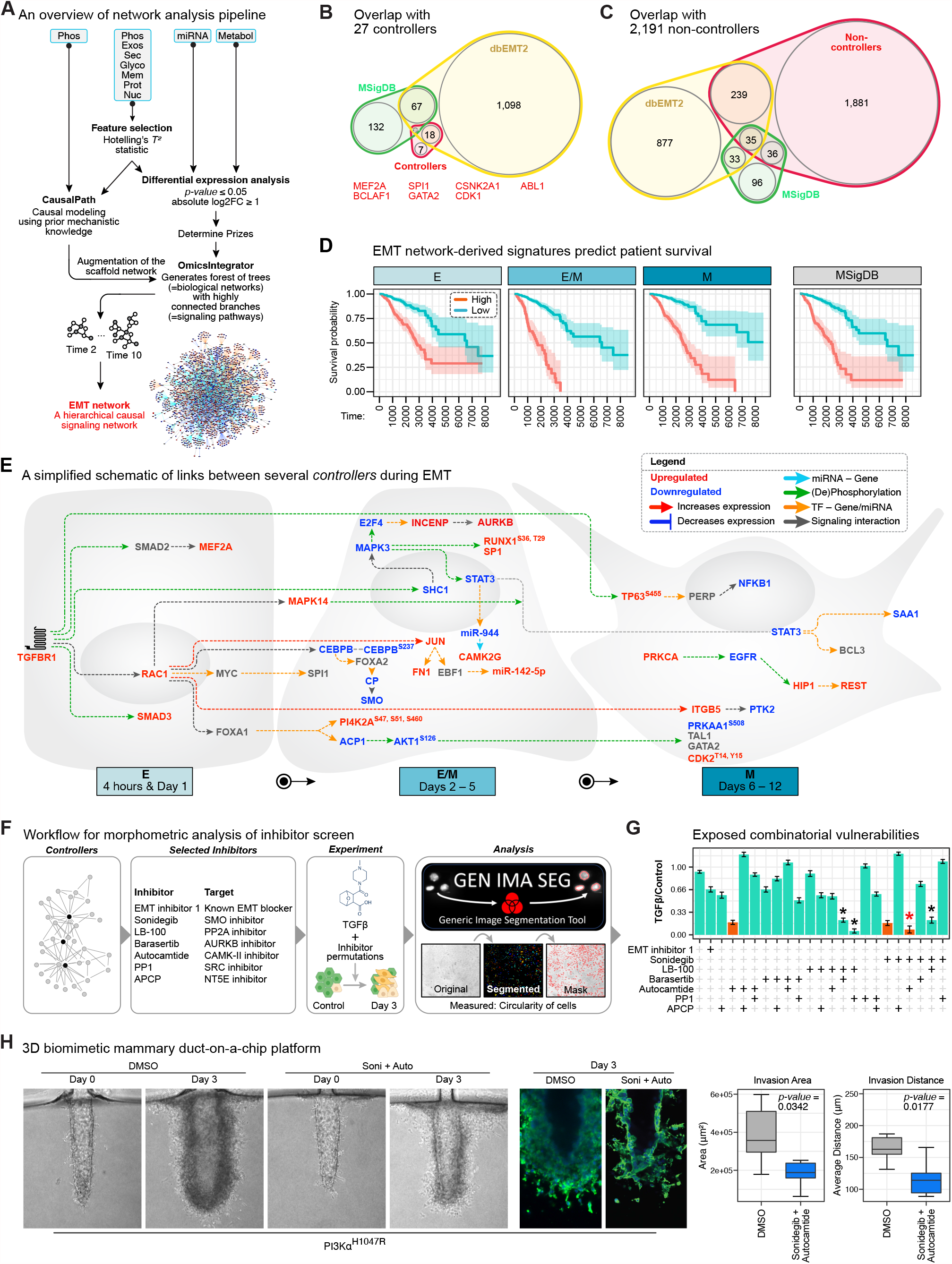
Integrative systems causal model of EMT. (A) Schematics of causal modeling workflow. Non-redundant genes with most significant expression profiles (Hotelling’s T^2^ statistic) were used. CausalPath-estimated logical networks were used to augment a custom-built confidence-weighted scaffold interactome which was then used to solve the Steiner Forest problem using the OmicsIntegrator software. Only differential molecules (relative to Control, FDR adj.*p-value* ≤ 0.05, |log2FC| ≥ 1) were considered as ‘prizes’. (B) Overlap between EMT databases and ‘controllers’ identified in this study. (C) Overlap between EMT databases and ‘non-controllers’ identified in this study. (D) Kaplan-Meier plots comparing prognostic performance of MSigDB hallmarks and ‘controllers’ identified in this study. (E) A simplified schematic showing the hierarchical relationships between several bottleneck genes at different stages of EMT, as indicated. (F) Workflow for morphometric screening of drug combinations. (G) Barplot of results of above experiment indicates varying degrees of synergy or antagonism between inhibitors, in influencing EMT-associated changes in cell shape (eccentricity). Error bars indicate spread of datapoints across all quantified cells in each condition. (H) 3D ducts were seeded with stable MCF10A^PIK3CA-H1047R^ cells and treated with Sonidegib and Autocamtide for 3 days. Area of invading cells and average distance traveled away from the ducts as compared to DMSO treated controls were quantified using ImageJ (n=6 devices).

One of the potential applications of the EMT network is to discover signaling paths from TGFBR1/2 to any gene(s) of interest within the network. As a demonstration, we queried several EMT-associated genes (FN1, MMP7, CD44, SCRIB, TWISTNB, ZEB1, SNAI2) and recovered previously known and unknown paths to them putatively active at multiple stages of EMT (**Fig. S7A**). Next, to identify key factors driving EMT, we performed ‘controllability’ analysis (Vinayagam et al., 2016) to identify controllers (nodes) exerting a significant influence on EMT network topology. Not surprisingly, a few of them are established key regulators of metastasis (**Fig. 7B**). Unlike controllers, however, the non-controller nodes were poorly represented in EMT literature (**Fig. 7C**), again highlighting gaps, and potentially identifying new regulatory processes. Survival analysis against a large publicly available dataset of primary breast cancers with long-term patient outcomes (Cancer Genome Atlas Network, 2012) showed a significant association between tumors with altered expression of these controllers and shortened overall survival (**Fig. 7D**).

We queried the EMT network to identify signaling contexts in which these controllers are active at various key stages of EMT, which could also provide clues into mechanistic vulnerabilities (**Fig. 7E**). As cells are stimulated with TGFβ, the TFs SMAD2 and SMAD3 are activated, as expected. Another early responder was RHO GTPase RAC1, an effector of both KRAS (Wu et al., 2014) and TGFβ signaling (Ungefroren et al., 2018), suggesting potential crosstalk. The downstream effector of RAC1, MAPK14 (p38 MAPK), was also regulated early in EMT, suggesting cooperation between RAC1 and MAPK pathways (Santibáñez et al., 2010). Our model suggests SMAD3 regulates two other TF hubs, CEBPB (CCAAT/enhancer-binding protein β) and FOXA1. Loss of CEBPB reportedly switches TGFβ signaling from growth-inhibiting to EMT-inducing (Johansson et al., 2013), while FOXA1 is reportedly a key TF during EMT (Wang et al., 2013). We observed that STAT3 is suppressed at later stages of EMT. A recent study in KRAS-driven lung and pancreatic cancer found that STAT3 is required for maintaining the E state and is lost during acquisition of M phenotypes (D’Amico et al., 2018).

The EMT network directly predicts novel avenues for blocking EMT. To assess this, we performed a morphometry-based screening where we treated MCF10A cells with TGFβ in combination with drugs which were predicted to inhibit several of the controllers active at E→E/M transition (**Fig. 7F**, see **STAR Methods**). Our analysis using a custom-built (Ochs et al., 2019) and publicly accessible image analysis software GENIMASEG (**Fig. 7F**) showed significant efficacy of LB100+Barasertib, LB100+PP1 and Sonidegib+Autocamtide in reverting the elongated phenotype of EMT-induced cells (**Fig. 7G**), thus providing direct experimental evidence to our predictions. Using a biomimetic 3D mammary duct-on-a-chip platform (Kutys et al., 2020), we further observed that combinatorically inhibiting SMO and CAMK-II (Sonidegib+Autocamtide) inhibits invasion driven by the PI3K variant, PIK3Cα^H1047R^, which is associated with chemo-refractoriness in a subset of triple negative BrCa patients (Janku et al., 2013).

Overall, our EMT network recapitulates known signaling pathways, uncovers novel routes of information flow to known regulators of EMT, identifies new signaling players and pathways, makes substantial and provocative novel predictions and reveals cohesive time-resolved regulatory patterns and mechanistic links between both controllers and non-controllers.

## DISCUSSION

EMT and cancer are *emergent systems* (Sigston and Williams, 2017) wherein progression through various stages is regulated by intricate networks of intra-and extra-cellular signaling within and between cells. The key to understanding such complex biological phenomena are establishing experimental workflows that integrate multiple tiers of biological information (Karczewski and Snyder, 2018).

Discussions on EMT are often guided by the Waddington metaphor of a ball (=cell) rolling over a phenotypic landscape, which is dynamically shaped by multiple parameters: topologies of signaling networks, molecular stochasticity, extraneous cooperating and opposing forces (e.g., EMT inducing and/or inhibitory ligands, interaction with other cells) (Li and Balazsi, 2018). By integrating several molecular layers, SOMs and ‘neighbor joining’ approaches revealed the kinetics of cell-fate transitions and major phenotypic switch points driven by TGFβ. Several studies have indicated molecular and phenotypic granularity in EMT continuum and suggested existence of discrete metastable E/M (Sha et al., 2019). Many current EMT markers are biased toward the later stages of EMT, when the process is approaching completion (Song et al., 2019). Consequently, the molecular nature of E/M is still poorly understood. Our time-course integrative SOM allowed us to trace the temporal unfolding and uncover the molecular nature of the E→M transition.

While previous omics studies have measured either one or two molecular layers to describe EMT, our multi-tiered datasets enabled the discovery of several new aspects of EMT which were not captured by previous approaches. For example, the correlation between *mRNA* and proteins were weak. Strikingly, correlations between the various *proteomics layers* were also found to be quite poor, indicating that systems behavior cannot readily be extrapolated by any single layer (e.g., mRNA or total proteome), but instead needs an integrated analysis of several molecular layers. We show that TGFβ-induced EMT is only partially driven by transcription, where several E genes are repressed only post-transcriptionally, while transcripts of M genes are upregulated but mostly during earlier time points. At later stages, post-translational mechanisms become more prominent in driving the process, suggesting that the regulatory control of EMT may be more flexible than previously appreciated. Similarly, our results also predict the mechanistic importance of protein subcellular localization during EMT. A comparison of proteins detected in *EV, Sec, Glyco, Mem* and *Nuc* indicated extensive regulation of protein localizations. Further, previous studies on EMT have largely focused on cell-autonomous signaling, whereas multiple inter-cellular signaling mechanisms are evident from our integrative analysis. Indeed, we found both *EV* and *Sec* to be extensively regulated during EMT. Once considered as cellular ‘garbage bins’, their active participation in signaling and crosstalk is increasingly recognized (H. Rashed et al., 2017). Our results demonstrating limited qualitative overlap between *EV* and *Sec* hint at a potentially overlooked mechanistic distinction and provides opportunities for new biological insights. We utilized our *Mem, Glyco* and *Sec* datasets to provide a repertoire of 79 putative new L-R pairs with potential roles during EMT, which should be validated in future studies. By combining these results with the scRNAseq profiles, we were able to define an extensive network of inter-cellular communications.

In conclusion, we have established a comprehensive multi-tiered molecular landscape of TGFβ-induced EMT. This study aims to provide a valuable resource which is accessible through an interactive website (https://www.bu.edu/dbin/cnsb/emtapp/) (**Fig. S7**) and will strongly complement hypothesis-driven research with direct implications for epithelial cancers.

## Author contributions

**Indranil Paul**: Conceptualization, Methodology, Software, Data curation, Writing – Original draft, Visualization, Investigation, Formal analysis

**Dante Bolzan**: Software, Formal analysis

**Ahmed Youssef**: Investigation, Software

**Keith A Gagnon**: Investigation

**Heather Joan Hook**: Investigation

**Gopal Karemore**: Software

**Michael Uretz John Oliphant**: Investigation

**Weiwei Lin**: Investigation

**Qian Liu**: Formal analysis

**Sadhna Phanse**: Software

**Dzmitry Padhorny**: Software

**Sergei Kotelnikov**: Software

**Carl White**: Software

**Guillaume P. Andrieu**: Investigation

**Christopher S. Chen**: Investigation

**Pingzhao Hu**: Formal analysis

**Gerald Denis**: Investigation

**Dima Kozakov**: Software

**Brian Raught**: Investigation

**Trevor Siggers**: Investigation

**Stefan Wuchty**: Software, Formal analysis

**Senthil Muthuswamy**: Conceptualization, Investigation

**Andrew Emili**: Conceptualization, Supervision, Writing – Original draft, Resources, Funding acquisition, Formal analysis

## Acknowledgements

We dedicate our sincere gratitude to Stefano Monti, Xarelabos Varelas, Valentina Perissi and Matthew Layne for providing critical feedback on the manuscript. We thank the Microarray and Sequencing Core Facility at Boston University School of Medicine for acquisition of mRNA, miRNA and scRNAseq data.

## Supplementary figure legends

**Figure S1. Related to Figure 1.**
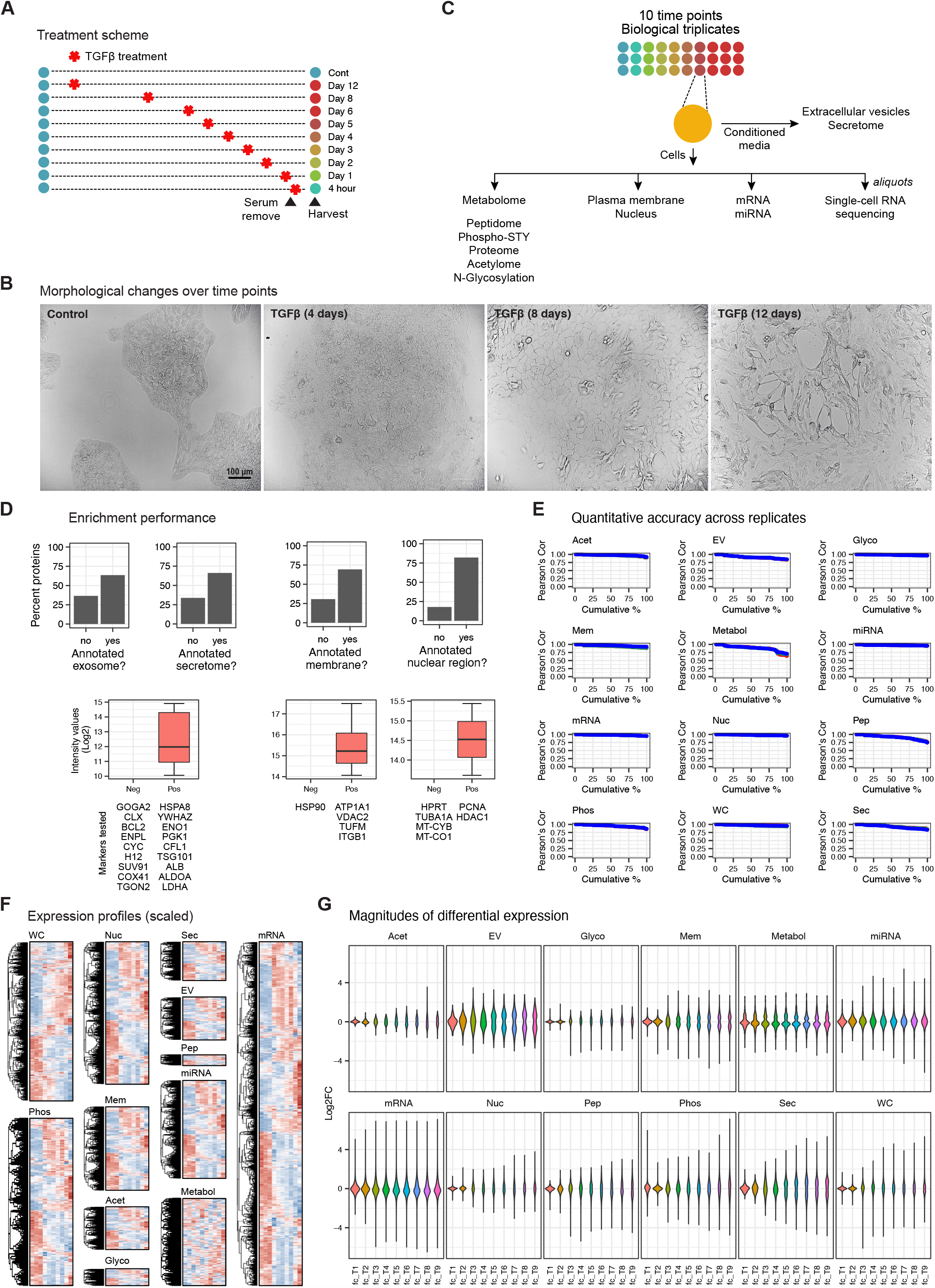
A multi-dimensional resource on TGFβ-induced EMT. (A) TGFβ treatments were staggered, at defined time periods, such that all plates were harvested at the same time. Cells were serum starved for 16 hours before harvesting. (B) Phase-contrast images of MCF10A cells at different time points. Scale bar = 100 µm. (C) For maximizing efficiency while maintaining compatibility with technology-specific protocols, after harvesting cells were collected in 4 aliquots per replicate, as shown. Conditioned media were also collected. (D) Upper panel – barplot showing percentage of quantified proteins with annotation in the ‘Cellular Component’ category. Lower panel – boxplot showing intensity values (log2) of proteins (=markers) commonly used to assess sub-cellular fractionation purity. Box edges correspond to 25th and 75th percentiles, whiskers include extreme data points. (E) Cumulative distribution (%) of Pearson’s coefficients across the samples (= time points). Significant overlap between the 3 biological replicates, shown as 3 colors, indicates high reproducibility. (F) Heatmaps of molecular expression profiles of indicated layers. The mean of all quantified molecules across the 3 replicates was used. (G) Violin plots showing the spread of log2FCs of molecules at each time point (relative to Control) for each layer.

**Figure S2. Related to Figure 2.**
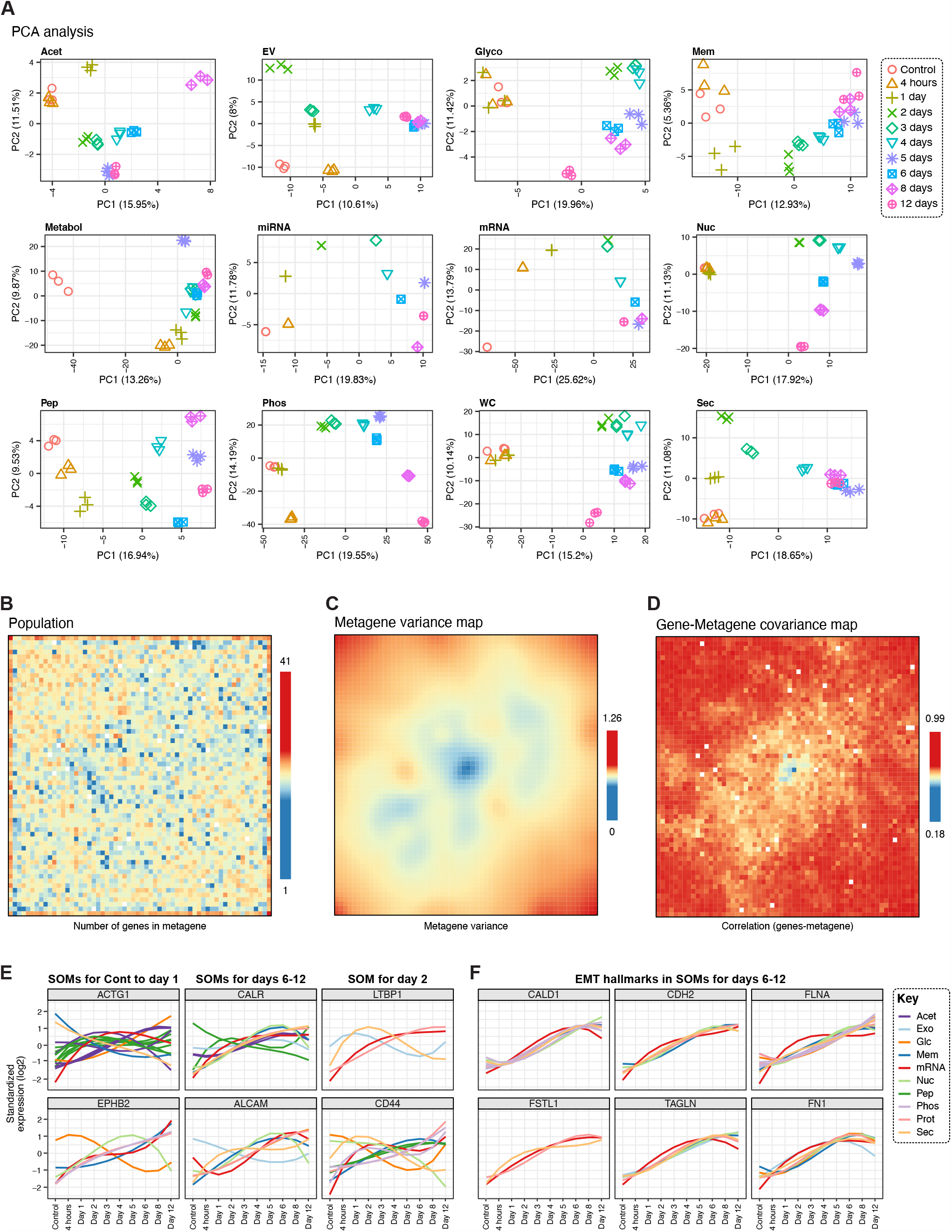
The topological architecture of EMT. (A) PCA of the various molecular layers. Time points are shown with different shapes and colors. Points with similar shape/color indicate biological replicates. (B) The population map presents the number of genes mapped to each individual metagene. (C) The plot summarizes co-variance structure of datasets at the metagene level. (D) The plot shows correlation of expression patterns of individual genes and the metagenes in which they are contained. (E) Temporal expression profiles of a few example genes identified in SOM analysis. Each colored line represents a molecular layer, as shown. (F) Temporal expression profiles of a few example genes enlisted as ‘EMT hallmarks’ in MSigDB database. Each colored line represents a molecular layer, as shown.

**Figure S3. Related to Figure 3.**
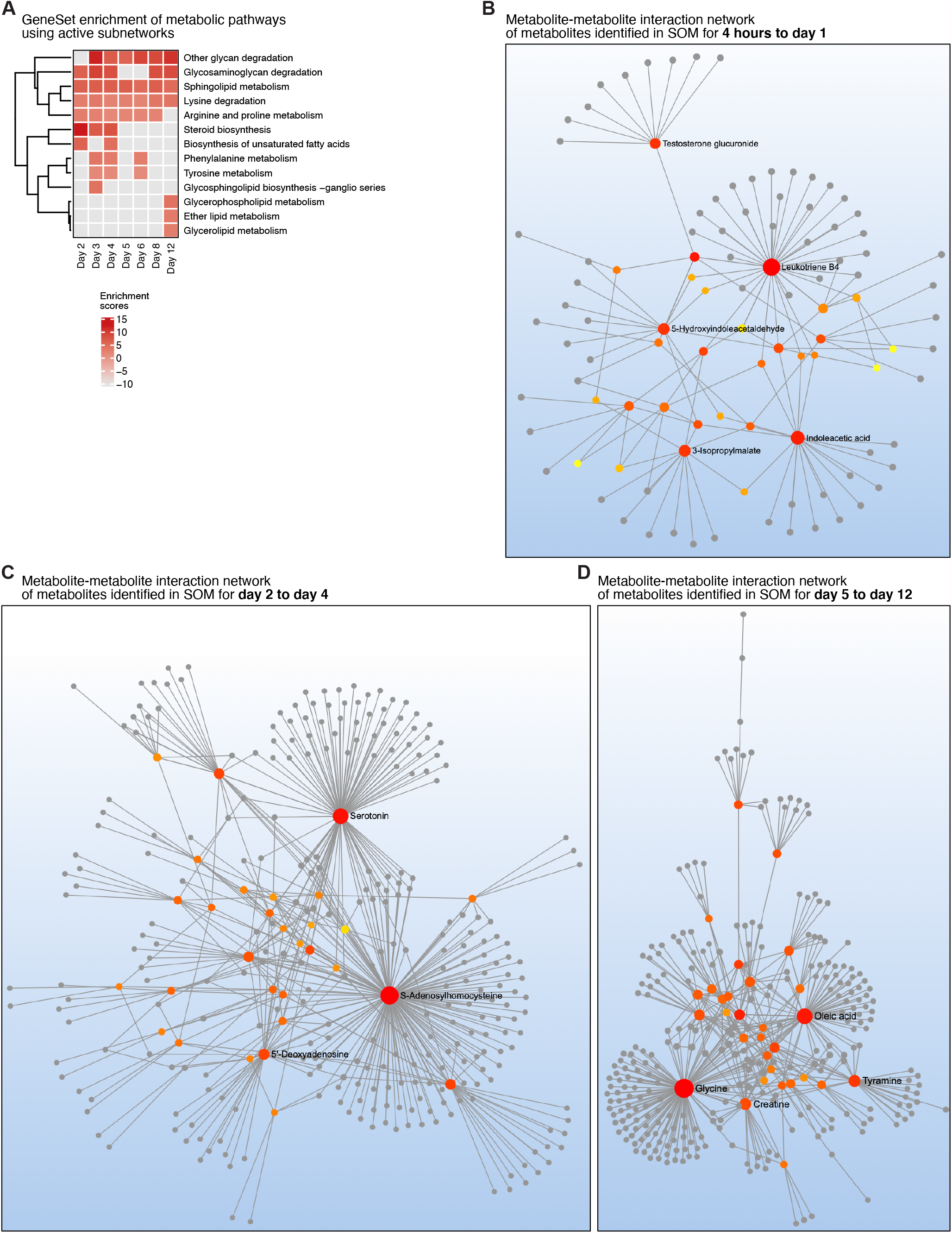
Dynamics of TGFβ-induced metabolic adaptations. (A) The heatmap displays the results of active subnetwork analysis using differential molecules in *WC* and *Phos* datasets. (B) – (D) Metabolite-metabolite interaction networks, using MetaboAnalyst, for indicated clusters. High connectivity indicates co-regulation of functionally related metabolites.

**Figure S4. Related to Figure 4.**
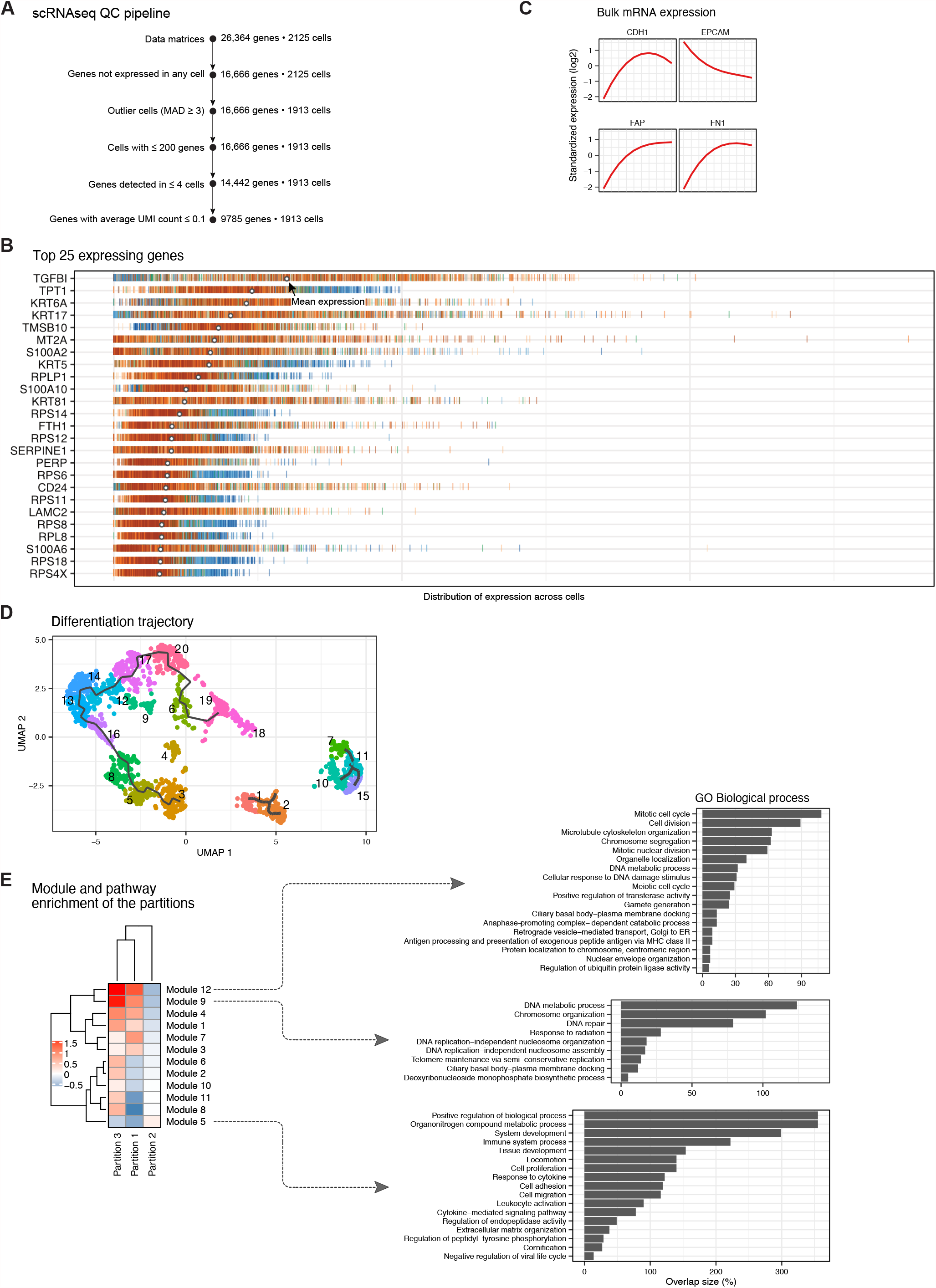
scRNAseq analysis reveals cellular dynamics and novel TFs for EMT. (A) An outline of the QC pipeline employed for scRNAseq data analysis. (B) The plot shows top 25 most expressed genes. Each row corresponds to a gene and each bar corresponds to the expression of the gene in single cells. (C) Expression of indicated genes in *mRNA* layer. (D) Developmental trajectories of MCF10A cells in response to TGFβ, inferred by Monocle3. Clusters are indicated by colors. (E) Heatmap showing aggregate expression of groups of genes (=Modules) with similar expression pattern across the *partitions*, by Monocle3. Modules 9/12 were highly expressed in P1/P3 and were enriched for ‘Cell cycle’ related GO annotations.

**Figure S5. Related to Figure 5.**
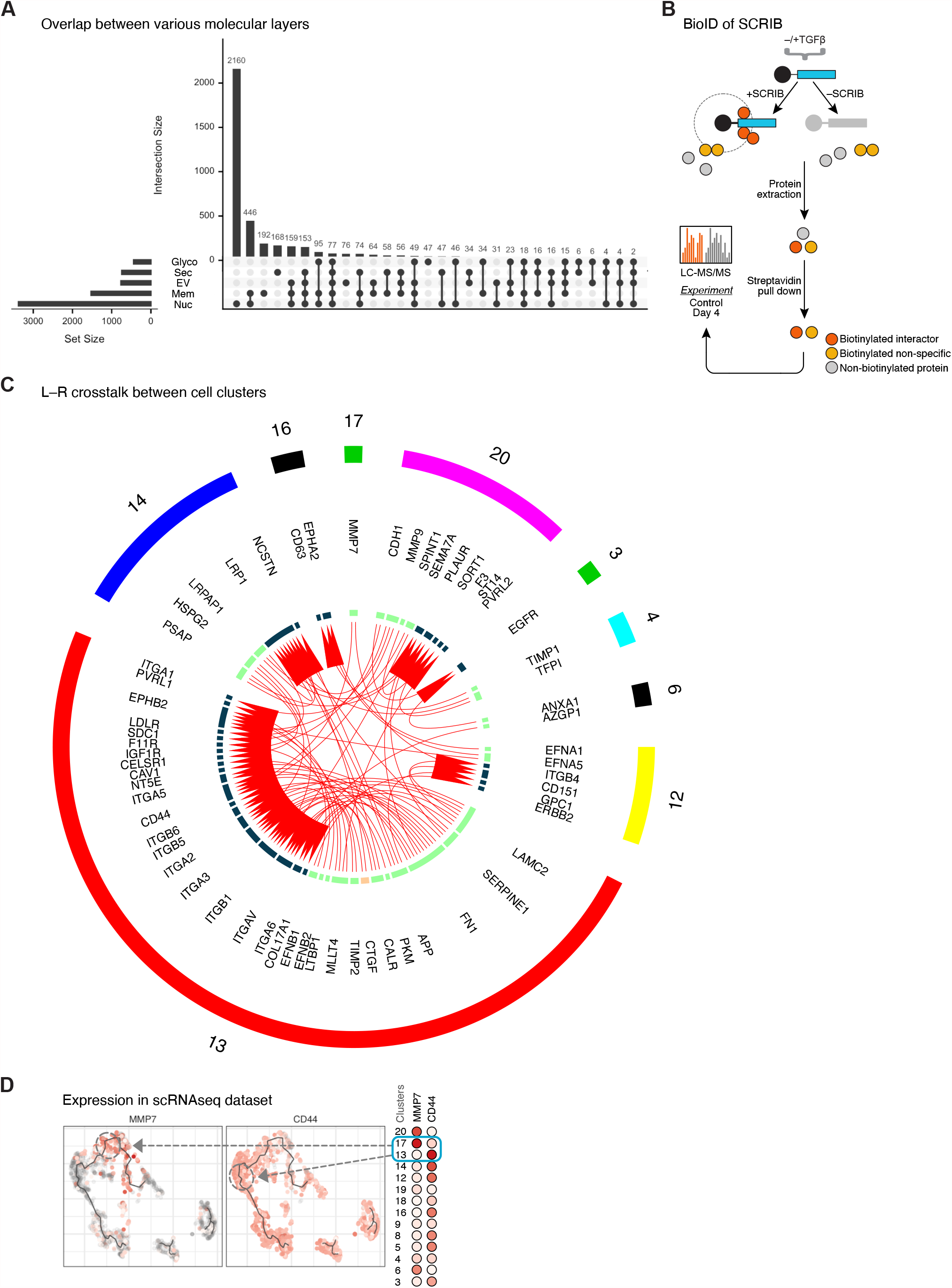
Spatial regulation of proteins and intercellular communication. (A) The plot shows number of common genes (intersection size, *y* axis) between layers as indicated. Only differential genes in each layer (‘Set size’, FDR ≤ 0.05; adj. *p-value* ≤ 0.05; r^2^ ≥ 0.6 & |log2FC|, relative to Control, of ≥ 1) were considered for the analysis. (B) Schematics of the BioID experiment performed to discover novel SCRIB partners induced by TGFβ signaling. (C) Circos plot showing L-R interactions between the P2 clusters. (D) UMAP plots of scRNAseq data highlighting the co-expression patterns of MMP7 and CD44.

**Figure S6. Related to Figure 6.**
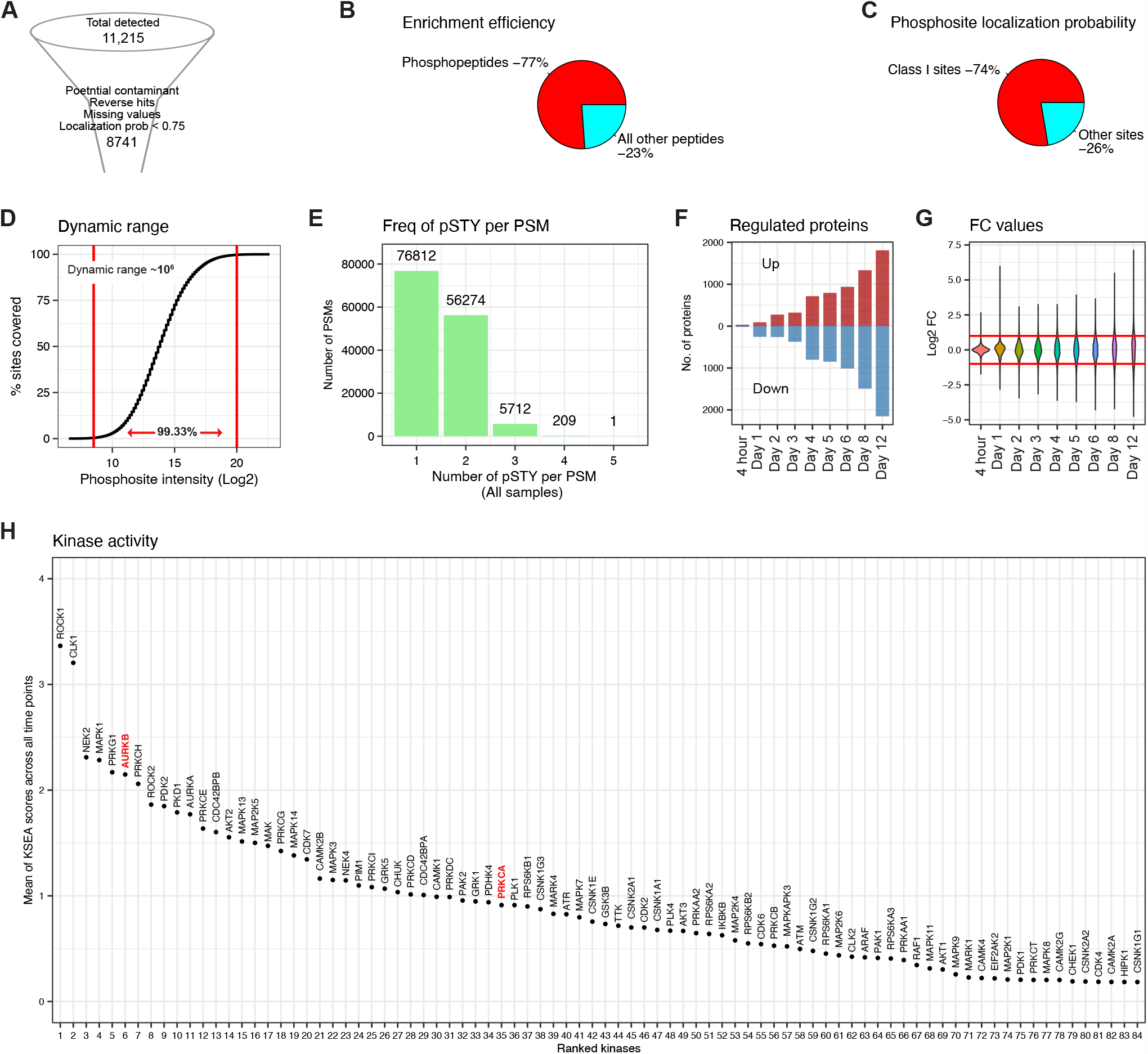
Phosphoproteome dynamics during EMT. (A) An outline of QC pipeline for *Phos* data analysis. (B) Enrichment efficiency for TiO_2_ workflow employed for the study. (C) About 74% of all detected p-sites were reliably localized (=Class I) by MaxQuant. (D) Quantifications of p-sites were achieved with a dynamic range of 10^6^ orders of magnitude. (E) Proportions of number of phosphate moieties in each detected p-site. (F) Number of phosphoproteins regulated over the time course. (G) Magnitudes of log2FC values for p-sites over the time course. (H) The plot shows all kinases ranked according to their KSEA enrichment. Mean of |z-score| values over all time points were taken.

**Figure S7. Related to Figure 7.**
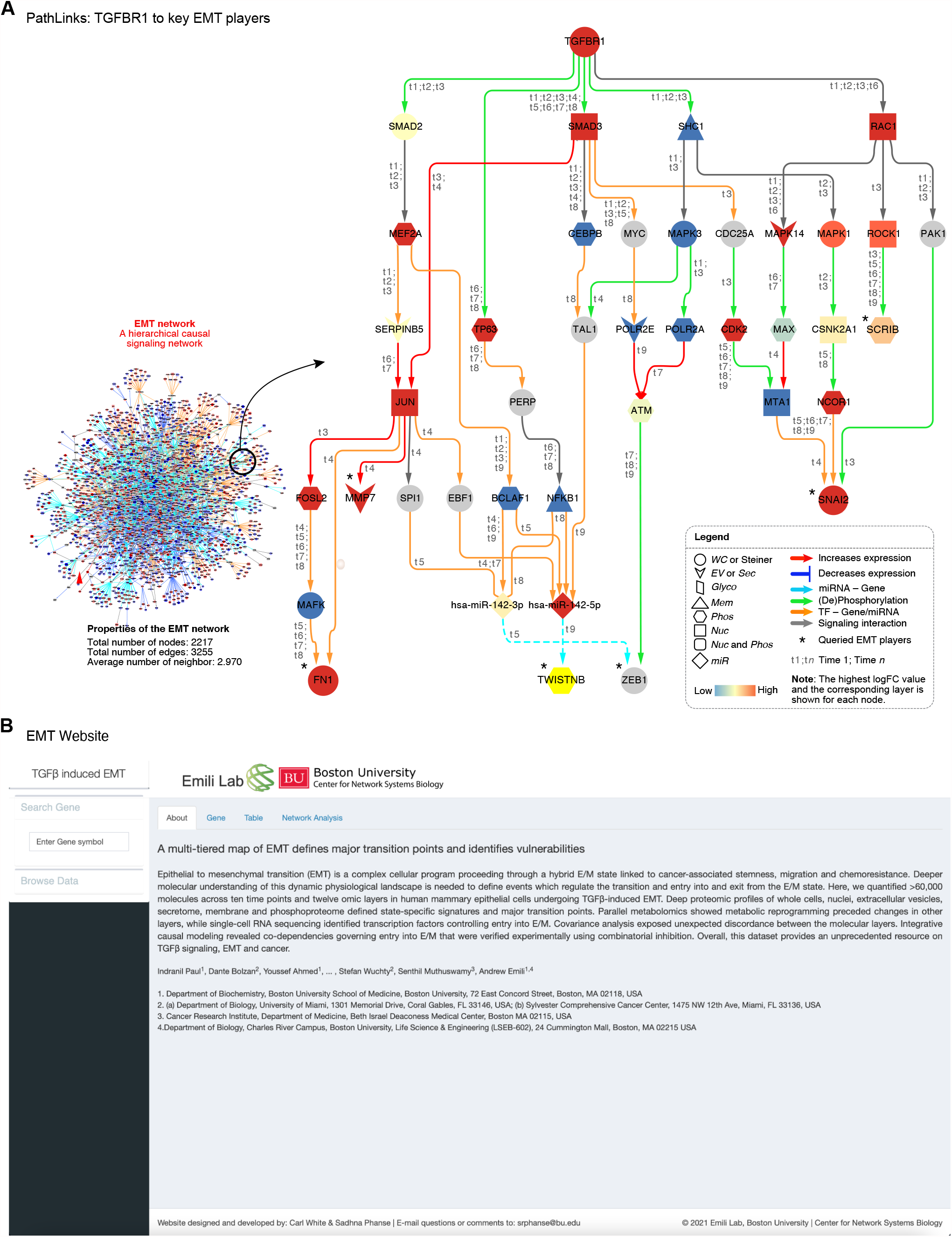
Integrative systems causal model of EMT. (A) A subgraph showing the signaling context of well-known EMT players ‘queried’ on the EMT network. (B) A snapshot of the companion website which can be interactively and freely accessed at https://www.bu.edu/dbin/cnsb/emtapp/.

## REFERENCES

Aibar, S., González-Blas, C.B., Moerman, T., Huynh-Thu, V.A., Imrichova, H., Hulselmans, G., Rambow, F., Marine, J.-C., Geurts, P., Aerts, J., et al. (2017). SCENIC: Single-cell regulatory network inference and clustering. Nat. Methods 14, 1083–1086.

Akhmedov, M., Kedaigle, A., Chong, R.E., Montemanni, R., Bertoni, F., Fraenkel, E., and Kwee, I. (2017). PCSF: An R-package for network-based interpretation of high-throughput data. PLOS Comput. Biol. 13, e1005694.

Bonello, T.T., and Peifer, M. (2019). Scribble: A master scaffold in polarity, adhesion, synaptogenesis, and proliferation. J. Cell Biol. 218, 742–756.

Cancer Genome Atlas Network (2012). Comprehensive molecular portraits of human breast tumours. Nature 490, 61–70.

Cerami, E., Gao, J., Dogrusoz, U., Gross, B.E., Sumer, S.O., Aksoy, B.A., Jacobsen, A., Byrne, C.J., Heuer, M.L., Larsson, E., et al. (2012). The cBio Cancer Genomics Portal: An Open Platform for Exploring Multidimensional Cancer Genomics Data: Figure 1. Cancer Discov. 2, 401–404.

Chin, Y.R., Yoshida, T., Marusyk, A., Beck, A.H., Polyak, K., and Toker, A. (2014). Targeting Akt3 Signaling in Triple-Negative Breast Cancer. Cancer Res. 74, 964–973.

Chong, J., Wishart, D.S., and Xia, J. (2019). Using MetaboAnalyst 4.0 for Comprehensive and Integrative Metabolomics Data Analysis. Curr. Protoc. Bioinforma. 68, e86.

Conesa, A., Nueda, M.J., Ferrer, A., and Talón, M. (2006). maSigPro: a method to identify significantly differential expression profiles in time-course microarray experiments. Bioinforma. Oxf. Engl. 22, 1096–1102.

D’Amico, S., Shi, J., Martin, B.L., Crawford, H.C., Petrenko, O., and Reich, N.C. (2018). STAT3 is a master regulator of epithelial identity and KRAS-driven tumorigenesis. Genes Dev. 32, 1175–1187.

Dongre, A., and Weinberg, R.A. (2019). New insights into the mechanisms of epithelial– mesenchymal transition and implications for cancer. Nat. Rev. Mol. Cell Biol. 20, 69–84.

D’Souza, R.C.J., Knittle, A.M., Nagaraj, N., Dinther, M. van, Choudhary, C., Dijke, P. ten, Mann, M., and Sharma, K. (2014). Time-resolved dissection of early phosphoproteome and ensuing proteome changes in response to TGF-β. Sci. Signal. 7, rs5–rs5.

Ferrell, J.E. (1998). How regulated protein translocation can produce switch-like responses. Trends Biochem. Sci. 23, 461–465.

Garg, M., Braunstein, G., and Koeffler, H.P. (2014). LAMC2 as a therapeutic target for cancers. Expert Opin. Ther. Targets 18, 979–982.

H. Rashed, M., Bayraktar, E., K. Helal, G., Abd-Ellah, M.F., Amero, P., Chavez-Reyes, A., and Rodriguez-Aguayo, C. (2017). Exosomes: From Garbage Bins to Promising Therapeutic Targets. Int. J. Mol. Sci. 18.

Hanna, V.S., and Hafez, E.A.A. (2018). Synopsis of arachidonic acid metabolism: A review. J. Adv. Res. 11, 23–32.

Hawe, J.S., Theis, F.J., and Heinig, M. (2019). Inferring Interaction Networks From Multi-Omics Data. Front. Genet. 10.

Heldin, C.-H., Vanlandewijck, M., and Moustakas, A. (2012). Regulation of EMT by TGFβ in cancer. FEBS Lett. 586, 1959–1970.

Hinz, N., and Jücker, M. (2019). Distinct functions of AKT isoforms in breast cancer: a comprehensive review. Cell Commun. Signal. CCS 17.

Hong, T., Watanabe, K., Ta, C.H., Villarreal-Ponce, A., Nie, Q., and Dai, X. (2015). An Ovol2-Zeb1 Mutual Inhibitory Circuit Governs Bidirectional and Multi-step Transition between Epithelial and Mesenchymal States. PLOS Comput. Biol. 11, e1004569.

Hua, W., ten Dijke, P., Kostidis, S., Giera, M., and Hornsveld, M. (2019). TGFβ-induced metabolic reprogramming during epithelial-to-mesenchymal transition in cancer. Cell. Mol. Life Sci.

Hughes, A.L., and Friedman, R. (2009). A phylogenetic approach to gene expression data: evidence for the evolutionary origin of mammalian leukocyte phenotypes. Evol. Dev. 11, 382–390.

Hung, M.-C., and Link, W. (2011). Protein localization in disease and therapy. J. Cell Sci. 124, 3381–3392.

Iliopoulos, D., Polytarchou, C., Hatziapostolou, M., Kottakis, F., Maroulakou, I.G., Struhl, K., and Tsichlis, P.N. (2009). MicroRNAs differentially regulated by Akt isoforms control EMT and stem cell renewal in cancer cells. Sci. Signal. 2, ra62.

Janku, F., Wheler, J.J., Naing, A., Falchook, G.S., Hong, D.S., Stepanek, V.M., Fu, S., Piha-Paul, S.A., Lee, J.J., Luthra, R., et al. (2013). PIK3CA mutation H1047R is associated with response to PI3K/AKT/mTOR signaling pathway inhibitors in early-phase clinical trials. Cancer Res. 73, 276–284.

Johansson, J., Berg, T., Kurzejamska, E., Pang, M.-F., Tabor, V., Jansson, M., Roswall, P., Pietras, K., Sund, M., Religa, P., et al. (2013). MiR-155-mediated loss of C/EBPβ shifts the TGF-β response from growth inhibition to epithelial-mesenchymal transition, invasion and metastasis in breast cancer. Oncogene 32, 5614–5624.

Karczewski, K.J., and Snyder, M.P. (2018). Integrative omics for health and disease. Nat. Rev. Genet. 19, 299–310.

Kim, E., Kim, J.-Y., Smith, M.A., Haura, E.B., and Anderson, A.R.A. (2018). Cell signaling heterogeneity is modulated by both cell-intrinsic and -extrinsic mechanisms: An integrated approach to understanding targeted therapy. PLoS Biol. 16.

Koundouros, N., and Poulogiannis, G. (2020). Reprogramming of fatty acid metabolism in cancer. Br. J. Cancer 122, 4–22.

Krishnaswamy, S., Zivanovic, N., Sharma, R., Pe’er, D., and Bodenmiller, B. (2018). Learning timevarying information flow from single-cell epithelial to mesenchymal transition data. PLoS ONE 13.

Kutys, M.L., Polacheck, W.J., Welch, M.K., Gagnon, K.A., Koorman, T., Kim, S., Li, L., McClatchey, A.I., and Chen, C.S. (2020). Uncovering mutation-specific morphogenic phenotypes and paracrinemediated vessel dysfunction in a biomimetic vascularized mammary duct platform. Nat. Commun. 11, 3377.

Lambert, A.W., Pattabiraman, D.R., and Weinberg, R.A. (2017). Emerging Biological Principles of Metastasis. Cell 168, 670–691.

Li, C., and Balazsi, G. (2018). A landscape view on the interplay between EMT and cancer metastasis. Npj Syst. Biol. Appl. 4, 1–9.

Li, C.-W., Xia, W., Lim, S.-O., Hsu, J.L., Huo, L., Wu, Y., Li, L.-Y., Lai, C.-C., Chang, S.-S., Hsu, Y.-H., et al. (2016). AKT1 Inhibits Epithelial-to-Mesenchymal Transition in Breast Cancer through Phosphorylation-Dependent Twist1 Degradation. Cancer Res. 76, 1451–1462.

Liu, X., Li, J., Cadilha, B.L., Markota, A., Voigt, C., Huang, Z., Lin, P.P., Wang, D.D., Dai, J., Kranz, G., et al. (2019). Epithelial-type systemic breast carcinoma cells with a restricted mesenchymal transition are a major source of metastasis. Sci. Adv. 5, eaav4275.

Liu, Y., Beyer, A., and Aebersold, R. (2016). On the Dependency of Cellular Protein Levels on mRNA Abundance. Cell 165, 535–550.

Ma, Y., Temkin, S.M., Hawkridge, A.M., Guo, C., Wang, W., Wang, X.-Y., and Fang, X. (2018). Fatty acid oxidation: An emerging facet of metabolic transformation in cancer. Cancer Lett. 435, 92–100.

McFaline-Figueroa, J.L., Hill, A.J., Qiu, X., Jackson, D., Shendure, J., and Trapnell, C. (2019). A pooled single-cell genetic screen identifies regulatory checkpoints in the continuum of the epithelial-to-mesenchymal transition. Nat. Genet. 51, 1389–1398.

Nieto, M.A., Huang, R.Y.-J., Jackson, R.A., and Thiery, J.P. (2016). EMT: 2016. Cell 166, 21–45.

Ochs, F., Karemore, G., Miron, E., Brown, J., Sedlackova, H., Rask, M.-B., Lampe, M., Buckle, V., Schermelleh, L., Lukas, J., et al. (2019). Stabilization of chromatin topology safeguards genome integrity. Nature 574, 571–574.

Pastushenko, I., Brisebarre, A., Sifrim, A., Fioramonti, M., Revenco, T., Boumahdi, S., Van Keymeulen, A., Brown, D., Moers, V., Lemaire, S., et al. (2018). Identification of the tumour transition states occurring during EMT. Nature 556, 463–468.

Pei, Y.-F., Liu, J., Cheng, J., Wu, W.-D., and Liu, X.-Q. (2019). Silencing of LAMC2 Reverses Epithelial-Mesenchymal Transition and Inhibits Angiogenesis in Cholangiocarcinoma via Inactivation of the Epidermal Growth Factor Receptor Signaling Pathway. Am. J. Pathol. 189, 1637–1653.

Pellinen, T., Blom, S., Sánchez, S., Välimäki, K., Mpindi, J.-P., Azegrouz, H., Strippoli, R., Nieto, R., Vitón, M., Palacios, I., et al. (2018). ITGB1-dependent upregulation of Caveolin-1 switches TGFβ signalling from tumour-suppressive to oncogenic in prostate cancer. Sci. Rep. 8, 1–14.

Puram, S.V., Parikh, A.S., and Tirosh, I. (2018). Single cell RNA-seq highlights a role for a partial EMT in head and neck cancer. Mol. Cell. Oncol. 5, e1448244.

Ramilowski, J.A., Goldberg, T., Harshbarger, J., Kloppmann, E., Lizio, M., Satagopam, V.P., Itoh, M., Kawaji, H., Carninci, P., Rost, B., et al. (2015). A draft network of ligand–receptor-mediated multicellular signalling in human. Nat. Commun. 6, 1–12.

Riemondy, K.A., Jansing, N.L., Jiang, P., Redente, E.F., Gillen, A.E., Fu, R., Miller, A.J., Spence, J.R., Gerber, A.N., Hesselberth, J.R., et al. (2019). Single-cell RNA sequencing identifies TGF-β as a key regenerative cue following LPS-induced lung injury. JCI Insight 4, e123637.

Sadej, R., Romanska, H., Kavanagh, D., Baldwin, G., Takahashi, T., Kalia, N., and Berditchevski, F. (2010). Tetraspanin CD151 Regulates Transforming Growth Factor β Signaling: Implication in Tumor Metastasis. Cancer Res. 70, 6059–6070.

Santibáñez, J.F., Kocić, J., Fabra, A., Cano, A., and Quintanilla, M. (2010). Rac1 modulates TGF-β1-mediated epithelial cell plasticity and MMP9 production in transformed keratinocytes. FEBS Lett. 584, 2305–2310.

Scheel, C., Eaton, E.N., Li, S.H.-J., Chaffer, C.L., Reinhardt, F., Kah, K.-J., Bell, G., Guo, W., Rubin, J., Richardson, A.L., et al. (2011). Paracrine and Autocrine Signals Induce and Maintain Mesenchymal and Stem Cell States in the Breast. Cell 145, 926–940.

Sha, Y., Haensel, D., Gutierrez, G., Du, H., Dai, X., and Nie, Q. (2019). Intermediate cell states in epithelial-to-mesenchymal transition. Phys. Biol. 16, 021001.

Shirakihara, T., Kawasaki, T., Fukagawa, A., Semba, K., Sakai, R., Miyazono, K., Miyazawa, K., and Saitoh, M. (2013). Identification of integrin α3 as a molecular marker of cells undergoing epithelial-mesenchymal transition and of cancer cells with aggressive phenotypes. Cancer Sci. 104, 1189–1197.

Sigston, E.A.W., and Williams, B.R.G. (2017). An Emergence Framework of Carcinogenesis. Front. Oncol. 7.

Song, J., Wang, W., Wang, Y., Qin, Y., Wang, Y., Zhou, J., Wang, X., Zhang, Y., and Wang, Q. (2019). Epithelial-mesenchymal transition markers screened in a cell-based model and validated in lung adenocarcinoma. BMC Cancer 19, 680.

Tabassum, D.P., and Polyak, K. (2015). Tumorigenesis: it takes a village. Nat. Rev. Cancer 15, 473–483.

Tallima, H., and El Ridi, R. (2018). Arachidonic acid: Physiological roles and potential health benefits – A review. J. Adv. Res. 11, 33–41.

Tam, W.L., Lu, H., Buikhuisen, J., Soh, B.S., Lim, E., Reinhardt, F., Wu, Z.J., Krall, J.A., Bierie, B., Guo, W., et al. (2013). Protein Kinase C α Is a Central Signaling Node and Therapeutic Target for Breast Cancer Stem Cells. Cancer Cell 24, 347–364.

Tamayo, P., Slonim, D., Mesirov, J., Zhu, Q., Kitareewan, S., Dmitrovsky, E., Lander, E.S., and Golub, T.R. (1999). Interpreting patterns of gene expression with self-organizing maps: Methods and application to hematopoietic differentiation. Proc. Natl. Acad. Sci. 96, 2907–2912.

Thomson, T.M., Balcells, C., and Cascante, M. (2019). Metabolic Plasticity and Epithelial-Mesenchymal Transition. J. Clin. Med. 8, 967.

Tominaga, K., Minato, H., Murayama, T., Sasahara, A., Nishimura, T., Kiyokawa, E., Kanauchi, H., Shimizu, S., Sato, A., Nishioka, K., et al. (2019). Semaphorin signaling via MICAL3 induces symmetric cell division to expand breast cancer stem-like cells. Proc. Natl. Acad. Sci. 116, 625–630.

Ulgen, E., Ozisik, O., and Sezerman, O.U. (2019). pathfindR: An R Package for Comprehensive Identification of Enriched Pathways in Omics Data Through Active Subnetworks. Front. Genet. 10.

Ungefroren, H., Witte, D., and Lehnert, H. (2018). The role of small GTPases of the Rho/Rac family in TGF-β-induced EMT and cell motility in cancer. Dev. Dyn. 247, 451–461.

Vinayagam, A., Gibson, T.E., Lee, H.-J., Yilmazel, B., Roesel, C., Hu, Y., Kwon, Y., Sharma, A., Liu, Y.-Y., Perrimon, N., et al. (2016). Controllability analysis of the directed human protein interaction network identifies disease genes and drug targets. Proc. Natl. Acad. Sci. U. S. A. 113, 4976–4981.

Wang, H., Meyer, C.A., Fei, T., Wang, G., Zhang, F., and Liu, X.S. (2013). A systematic approach identifies FOXA1 as a key factor in the loss of epithelial traits during the epithelial-to-mesenchymal transition in lung cancer. BMC Genomics 14, 680.

Wirth, H., von Bergen, M., and Binder, H. (2012). Mining SOM expression portraits: feature selection and integrating concepts of molecular function. BioData Min. 5, 18.

Wu, C.-Y.C., Carpenter, E.S., Takeuchi, K.K., Halbrook, C.J., Peverley, L.V., Bien, H., Hall, J.C., DelGiorno, K.E., Pal, D., Song, Y., et al. (2014). PI3K regulation of RAC1 is required for KRAS-induced pancreatic tumorigenesis in mice. Gastroenterology 147, 1405-1416.e7.

Yang, J., Antin, P., Berx, G., Blanpain, C., Brabletz, T., Bronner, M., Campbell, K., Cano, A., Casanova, J., Christofori, G., et al. (2020). Guidelines and definitions for research on epithelial–mesenchymal transition. Nat. Rev. Mol. Cell Biol. 1–12.

Zhang, J., Tian, X.-J., Zhang, H., Teng, Y., Li, R., Bai, F., Elankumaran, S., and Xing, J. (2014). TGF-β– induced epithelial-to-mesenchymal transition proceeds through stepwise activation of multiple feedback loops. Sci Signal 7, ra91–ra91.

Zhang, J., Lin, X., Wu, L., Huang, J.-J., Jiang, W.-Q., Kipps, T.J., and Zhang, S. (2020). Aurora B induces epithelial–mesenchymal transition by stabilizing Snail1 to promote basal-like breast cancer metastasis. Oncogene 39, 2550–2567.

Zhang, K., Myllymäki, S.-M., Gao, P., Devarajan, R., Kytölä, V., Nykter, M., Wei, G.-H., and Manninen, A. (2017). Oncogenic K-Ras upregulates ITGA6 expression via FOSL1 to induce anoikis resistance and synergizes with αV-Class integrins to promote EMT. Oncogene 36, 5681–5694.

